# MINT: a toolbox for the analysis of multivariate neural information coding and transmission

**DOI:** 10.1101/2024.11.04.621910

**Authors:** Gabriel Matías Lorenz, Nicola M. Engel, Marco Celotto, Loren Kocillari, Sebastiano Curreli, Tommaso Fellin, Stefano Panzeri

**Affiliations:** Institute for Neural Information Processing, Center for Molecular Neurobiology (ZMNH), University Medical Center Hamburg-Eppendorf (UKE), Hamburg, Germany; Istituto Italiano di Tecnologia, Genoa, Italy; Department of Pharmacy and Biotechnology, University of Bologna, Bologna, Italy; Department of Brain and Cognitive Sciences, Picower Institute for Learning and Memory, Massachusetts Institute of Technology, Cambridge, MA, USA; Department of Neurophysiology and Pathophysiology, University Medical Center Hamburg-Eppendorf (UKE), Hamburg, Germany

## Abstract

Information theory has deeply influenced the conceptualization of brain information processing and is a mainstream framework for analyzing how neural networks in the brain process information to generate behavior. Information theory tools have been initially conceived and used to study how information about sensory variables is encoded by the activity of small neural populations. However, recent multivariate information theoretic advances have enabled addressing how information is exchanged across areas and used to inform behavior. Moreover, its integration with dimensionality-reduction techniques has enabled addressing information encoding and communication by the activity of large neural populations or many brain areas, as recorded by multichannel activity measurements in functional imaging and electrophysiology. Here, we provide a Multivariate Information in Neuroscience Toolbox (MINT) that combines these new methods with statistical tools for robust estimation from limited-size empirical datasets. We demonstrate the capabilities of MINT by applying it to both simulated and real neural data recorded with electrophysiology or calcium imaging, but all MINT functions are equally applicable to other brain-activity measurement modalities. We highlight the synergistic opportunities that combining its methods afford for reverse engineering of specific information processing and flow between neural populations or areas, and for discovering how information processing functions emerge from interactions between neurons or areas. MINT works on Linux, Windows and macOS operating systems, is written in MATLAB (requires MATLAB version 2018b or newer) and depends on 5 native MATLAB toolboxes. The calculation of one possible way to compute information redundancy requires the installation and compilation of C files (made available by us also as pre-compiled files). MINT is freely available at https://github.com/panzerilab/MINT with DOI 10.5281/zenodo.13998526 and operates under a GNU GPLv3 license.

## Introduction

Brain functions are based on the ability of groups of neurons or brain areas to encode, process and transmit information [1, 2]. Consequently, information theory [3], the mathematical theory of communication, has deeply influenced the conceptualization of brain operations. It has become a method of choice to analyze neural activity because of its many advantages [4-8]. It provides single-trial measures of how neural activity encodes variables important for cognitive functions such as sensory stimuli, and it is thus more relevant for single-trial behavior than trial-averaged measures. It captures contributions of both linear and non-linear interactions between variables at all orders, and thus allows hypotheses-free measures of information encoding that place upper bounds to the performance of any decoder. Because of its generality, it can be applied to any type of brain activity recordings. Also, it facilitates direct comparisons between the predictions of normative neural theories and real neural data [6, 9].

Earlier work using information theory to analyze empirical neural data has focused on low-dimensional measures of neural activity such as single neurons, small neural populations or aggregate measures (LFPs, M/EEG, fMRI). These studies have considered only how information is encoded in neural activity, regardless of how it may be used downstream. Such seminal studies have demonstrated e.g. how the temporal structure of neural activity (from single-neuron spike timing to network oscillations [10-16]) contributes to sensory encoding, or how neural mechanisms such as adaptation contribute to brain information processing [8, 17].

Over the last decade, neuroscience has seen major progress in the ability to record simultaneously the activity of many neurons and/or brain areas. These advances have driven the development of novel information theoretic analytical tools to investigate how information processing emerges from the interaction and communication among neurons or areas. Studies have provided multivariate information tools to individuate when synergy and redundancy arise in small populations, or to understand the mechanisms for generating redundancy and synergy, for example to characterize how correlations between the activity of different neurons shape information processing [1, 18-22]. Recent work has also coupled information theory with dimensionality-reduction techniques to study how information is encoded in populations of tens to hundreds of cells [23-30]. Other studies have developed multivariate information theory to quantify transmission, rather than encoding, of information across neurons or areas [31-40]. These methods measure the overall or stimulus-specific information exchanged between simultaneously recorded neurons and areas, and determine whether transmission relies on synergistic integration of information across nodes. Another major direction of progress has been in recording neural activity during behavior [41]. To support the growing interest on how neural computations shape behavior, information theory has produced tools to characterize the multivariate simultaneous relationship between sensory stimuli, neural activity and behavioral output to enable quantifying the impact on behavior of the information encoded in a certain area or population [24, 42, 43].

While the use and dissemination of information theoretic algorithms has been aided by software toolboxes [44-59], no toolbox yet provides a comprehensive implementation of tools to compute both information encoding and transmission, to break down information into components reflecting the effect of interactions and to quantify behavioral or downstream relevance of the encoded information (see Table S1). To fill these gaps and address the need to collect organically these tools in a format that allows immediately multiple analyses, here we introduce a new Multivariate Information in Neuroscience Toolbox (MINT). MINT provides a comprehensive set of information theoretic functions (including Shannon Entropy and Mutual Information, directed information transmission measures, information decompositions) and estimators (binned probability estimators, limited-sampling bias corrections). The implemented information-theoretic functions are detailed in Supplemental Material. What they compute, and how they can be used in neuroscience is summarized in Table 1. The accuracy and applicability of these algorithms has been validated and demonstrated extensively with both discrete neural variables, such as spikes in electrophysiological recordings [8, 13, 14, 60-62], and continuous neural variables, e.g. LFP, M/EEG, fMRI and calcium traces [24, 30, 63-65].

**Table 1:**
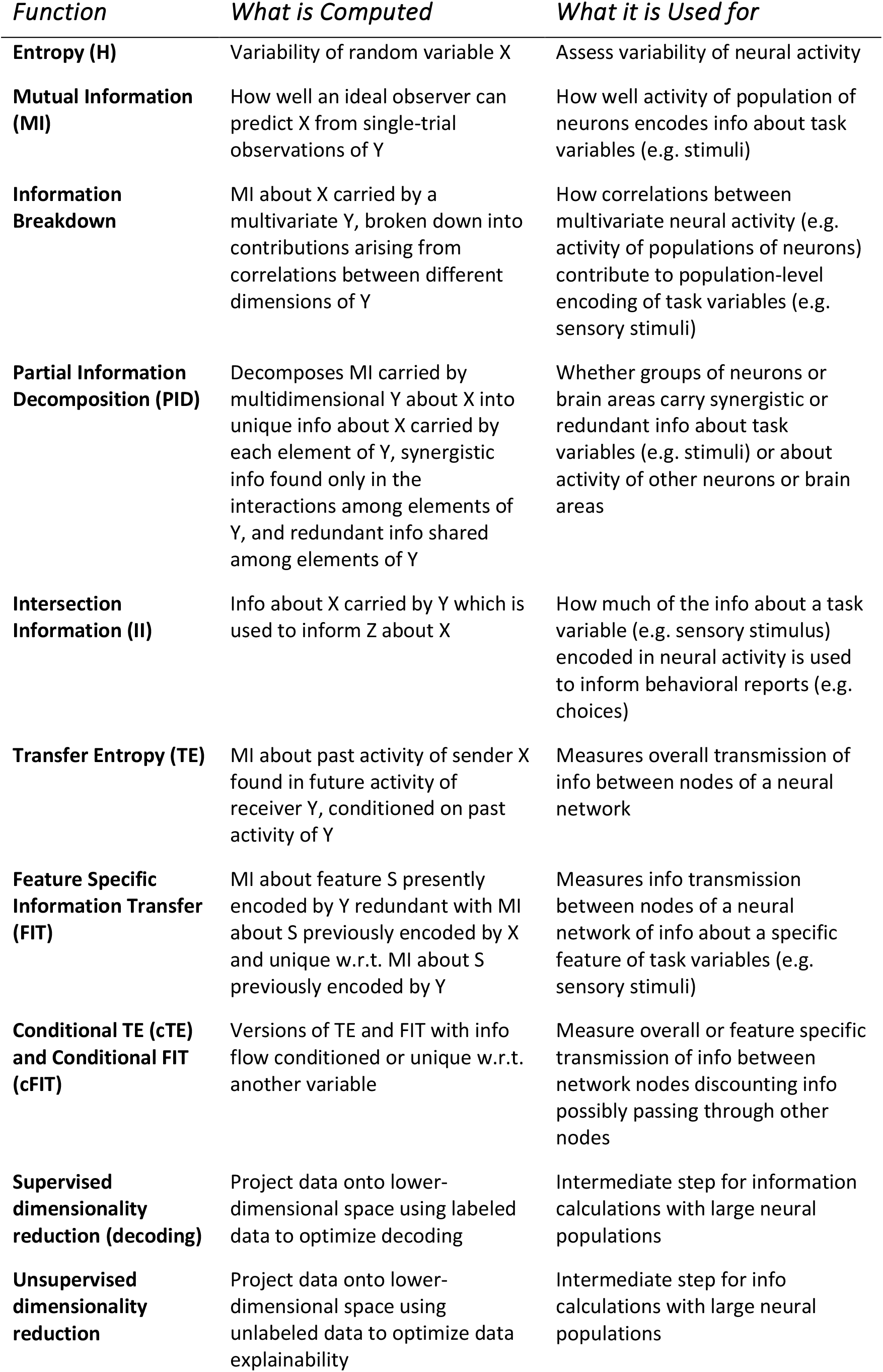
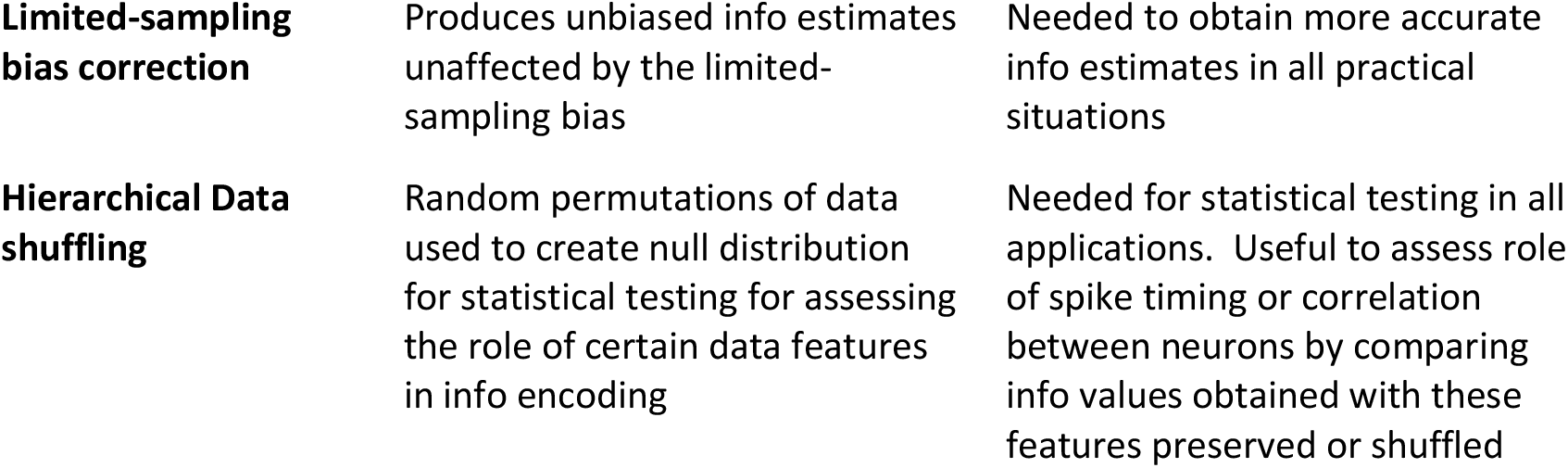
Glossary of Main Information Theoretic Functions. This table reports a short explanation of what the implemented information theoretic quantities
compute and for what type of applications they may be used. X, Y, Z denote random variables.

| *Function* | *What is Computed* | *What it is Used for* |
| --- | --- | --- |
| **Entropy (H)** | Variability of random variable X | Assess variability of neural activity |
| **Mutual Information (MI)** | How well an ideal observer can predict X from single-trial observations of Y | How well activity of population of neurons encodes info about task variables (e.g. stimuli) |
| **Information Breakdown** | MI about X carried by a multivariate Y, broken down into contributions arising from correlations between different dimensions of Y | How correlations between multivariate neural activity (e.g. activity of populations of neurons) contribute to population-level encoding of task variables (e.g. sensory stimuli) |
| **Partial Information Decomposition (PID)** | Decomposes MI carried by multidimensional Y about X into unique info about X carried by each element of Y, synergistic info found only in the interactions among elements of Y, and redundant info shared among elements of Y | Whether groups of neurons or brain areas carry synergistic or redundant info about task variables (e.g. stimuli) or about activity of other neurons or brain areas |
| **Intersection Information (II)** | Info about X carried by Y which is used to inform Z about X | How much of the info about a task variable (e.g. sensory stimulus) encoded in neural activity is used to inform behavioral reports (e.g. choices) |
| **Transfer Entropy (TE)** | MI about past activity of sender X found in future activity of receiver Y, conditioned on past activity of Y | Measures overall transmission of info between nodes of a neural network |
| **Feature Specific Information Transfer (FIT)** | MI about feature S presently encoded by Y redundant with MI about S previously encoded by X and unique w.r.t. MI about S previously encoded by Y | Measures info transmission between nodes of a neural network of info about a specific feature of task variables (e.g. sensory stimuli) |
| **Conditional TE (cTE) and Conditional FIT (cFIT)** | Versions of TE and FIT with info flow conditioned or unique w.r.t. another variable | Measure overall or feature specific transmission of info between network nodes discounting info possibly passing through other nodes |
| **Supervised dimensionality reduction (decoding)** | Project data onto lower-dimensional space using labeled data to optimize decoding | Intermediate step for information calculations with large neural populations |
| **Unsupervised dimensionality reduction** | Project data onto lower-dimensional space using unlabeled data to optimize data explainability | Intermediate step for info calculations with large neural populations |
| **Limited-sampling bias correction** | Produces unbiased info estimates unaffected by the limited-sampling bias | Needed to obtain more accurate info estimates in all practical situations |
| **Hierarchical Data shuffling** | Random permutations of data used to create null distribution for statistical testing for assessing the role of certain data features in info encoding | Needed for statistical testing in all applications. Useful to assess role of spike timing or correlation between neurons by comparing info values obtained with these features preserved or shuffled |

Importantly, as we demonstrate with examples, combining these multivariate tools enables addressing questions that cannot be addressed with a single tool. For example, combining tools to identify the specific contribution of correlations to population encoding or the amount of encoded information that informs behavior with dimensionality-reduction techniques allows understanding how large neural populations influence behaviors. Combining information encoding tools with content-specific information transmission tools can reverse engineer information flow in neural networks with unprecedented understanding. We thus anticipate that MINT will lead to uncover numerous new insights into neural information processing.

## Design and implementation

MINT is written in MATLAB (version 2018b or newer) and depends on the Statistics and Machine Learning, Optimization, Parallel Computing, Communications, and DSP System Toolboxes. MINT takes as input neural data (array of neural activity recorded in each trial) and task variables (sensory stimuli or behavioral responses presented or produced in each trial). It outputs information values and their null-hypothesis values for computing statistical significance. Fig. S1 illustrates MINT functions, options, and core routines.

MINT computes Entropy (`H.m`), which measures neural variability; Mutual Information (`MI.m`), which measures information encoding (Fig. 1B); (Fig. 1D). It computes the Information Breakdown of Shannon Information into contributions due to correlations between neurons [18, 20, 21, 66, 67]. It also computes Partial Information Decompositions (PID) [68, 69] of the information about a target variable carried by two or more source variables into unique, synergistic and redundant information (Fig. 1C). Computation of PID requires specifying a redundancy measure, which can be selected by the user from major options [52, 68, 70, 71] with complementary advantages. (Redundancy of [52] requires either a MATLAB-compatible C compiler or pre-compiled files made available by us for Windows 11, macOS, and Linux Debian). MINT computes additional functions of neuroscientific value: Intersection Information (II, function `II.m`; Fig. 1A), the amount of stimulus information in neural activity that is used to inform behavior; Transfer Entropy [72] (TE) and Feature-Specific Information Transfer (FIT), which measure overall and stimulus-feature-specific information transmission between nodes of neural networks (`TE.m`, `FIT.m` and `cFIT.m`; Fig. 1D).

**Figure 1:**
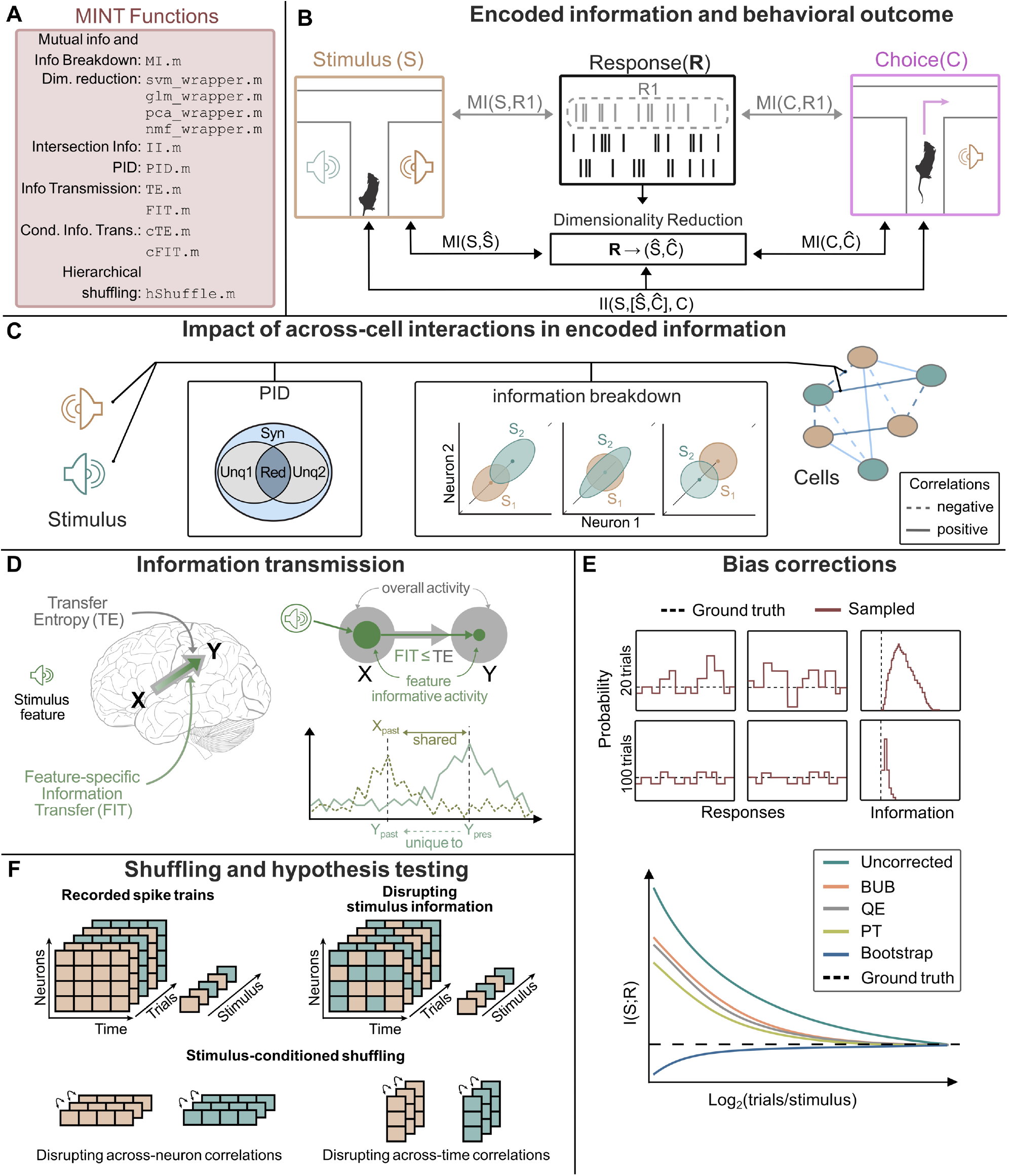
Overview of MINT. **A:** List of main MINT functions. **B:** MINT provides multivariate information theoretic functions to quantify the amount of information that single neurons or neural populations carry about task-relevant variables (e.g. sensory stimuli or behavioral choices). These methods are based on either direct calculation of information from maximum likelihood estimators of the discretized probabilities (ideal for small neural populations but not scalable with population size) or by using supervised or unsupervised dimensionality-reductions techniques to approximate high-dimensional neural population response probabilities with probabilities in lower-dimensional spaces (scalable with population size). It also provides tools to quantify how much of the information encoded by neural activity is used to inform behavior. **C:** MINT has multiple functions to compute, in small or large populations, how interactions between the activity of different neurons shape information encoding and create synergy or redundancy. **D:** MINT has tools to compute transfer of information from one neural population or brain region to another. It can compute both the total or stimulus-specific information transmitted between two nodes, with the option of conditioning over the activity of other nodes. **E:** MINT has tools to correct for the limited sampling bias that plague information estimates from limited datasets, an essential tool for analysis of empirical neuroscience data. **F:** MINT has a set of hierarchical permutation algorithms that provide null hypothesis testing for significance of information encoding and information transmission and for the impact of correlations across neurons or time.

The information quantities depend on the probabilities of task variables (e.g. presented sensory stimuli) and neural responses. MINT implements maximum likelihood probability estimators based on discretizing neural responses and task variable values and computing the empirical occurrences across experimental trials. These estimators have been widely used in neuroscience information theoretic studies, because neural spiking activity is intrinsically discrete and is usually quantified as the number of spikes emitted in one-or multiple-time windows of interest [14, 73]. Because they are simple and do not make assumptions about the probability distributions, these discretized estimators have been used to compute information also from continuous-valued aggregate measures of neural activity such as LFP, M/EEG, fMRI [11, 63, 64, 74] or continuous-valued behavioral variables [75]. MINT provides binning functions to discretize analogue data (equi-spaced or equi-populated binning, binning with user-defined bin edges, and possibly automated determination of bin numbers [76, 77]).

Any real experiment only yields a finite number of trials from which probabilities must be estimated. Finite sampling leads to a systematic error (bias) in information estimates (Fig. 1E), which can be as big as the true information values. Thus, bias corrections methods are essential for practical neuroscience applications, and six such well-established methods are included in MINT [78-82]. These methods, along with binning, parallelization options and other features are user-specified in an input structure (`opts`).

Information is computed by default with the so called *direct method* [14, 73], which computes information by directly computing probabilities as empirically occurrences of discrete or binned responses (function `MI.m`). The advantage of this method is that it preserves all information available in the discretized neural activity. For small-dimensional (up to N=2 or 3) neural response (e.g. responses of populations of up to 2-3 neurons) it is the recommended method as its estimates from datasets of realistic sizes can be still effectively corrected for the limited-sampling bias (Fig. S3). When considering high-dimensional neuronal responses (such as the activity of populations of many neurons) the curse of dimensionality prevents the direct sampling of the joint response probabilities from the data (Fig. S3).

We thus provide additional pipelines, recommended for high-dimensional neural responses such as the activity of large neural populations, that compute information from the empirical neural response probabilities but after reducing the dimensionality of neural population activity[24]. These *dimensionality-reduction pipelines* include supervised methods (Support Vector Machines, SVMs and Generalized Linear Models, GLMs) which reduce the dimensionality by providing decoding or posterior probabilities of the task variables given the single-trial neural population activity (Fig. 1A) and allow reliable estimations with small datasets (Fig. S3). We also provide unsupervised methods (Non-negative Matrix Factorization, NMF [83]; Principal Component Analysis, PCA) which reduce dimensionality individuating small numbers of dimensions with the highest explanation power of neural activity. MINT provides all these dimensionality-reduction techniques with native MATLAB functions, but it also allows easy interfacing with external libraries (e.g. libsvm [84] and glmnet [85]) (Fig. S2). Importantly, these dimensionality-reduction tools can be coupled with MINT’s *Hierarchical Shuffling* tools which can disrupt, by trial shuffling, specific features of population activity (such as response timing or correlations between neurons) to probe their contribution to information processing [24, 86].

## Results

We illustrate how to use MINT to address highly topical neuroscientific questions, emphasizing the utility of using synergistically multiple algorithms, allowed by MINT. In all examples, we use the limited-sampling bias corrections and hierarchical data shuffles of MINT, as they are essential for empirical data analyses.

### Computing the role of interactions between neurons in information encoding

An important question in neuroscience is whether and how the functional interactions (measured as activity correlations) between neurons enhance or limit information encoding in neural populations [1, 87]. Several information theoretic methods have been developed to address complementary aspects of this question [18-22, 66, 68, 86, 88]. Here we illustrate what we gain from their combined usage enabled by MINT.

We consider how a population of N neurons encodes information about a stimulus variable S. For neuron pairs (N=2), we computed the population information (Mutual Information between stimulus and the joint neural population response) with the direct method that estimates information directly form the empirical discretized response probabilities (see Design and implementation). The overall effect of interactions between neurons is expressed by the co-information, the difference between the population information and the sum of single-neuron stimulus information [19]. Positive (negative) co-information indicates predominantly synergistic (redundant) interactions. Contributions of synergy and redundancy to co-information can be separated using PID [69, 89]. The Information Breakdown [20-22] shows how co-information arises from interactions between neurons, by breaking down co-information into components *I*_*sig*−*sim*_ (contribution of the similarity across neurons of trial-averaged responses to different stimuli, see also [19]), *I*_*cor*−*ind*_ (contribution of the interplay between the signs of signal similarity and of noise correlations, defined as correlations between neurons in trials to the same stimulus), and of *I*_*cor*−*dep*_ (quantifying information added by the stimulus-modulation of noise correlations, or, equivalently, bounding the information lost when using decoders trained without considering correlations [18, 67]).

These small-population direct information calculations have the advantage of not making assumptions about decoding mechanisms, but do not scale up to large populations because of the curse of dimensionality [80]. Population information can be obtained by estimating probabilities in the reduced space of the stimuli decoded from single-trial neural activity. These estimates scale well with population size and can be computed robustly with small datasets (Fig. S3). However, they may underestimate total information in neural activity (see Fig. 3A), as direct-method information bound from above the information that can be extracted by any decoder, and they depend on the specific form of the considered decoder. We illustrate below how MINT’s decoding and data-shuffling tools allow to manipulate whether or not decoders consider information in correlated activity to gain information about the encoding mechanisms [1, 24, 86].

We illustrate these methods first by simulating the activity of N=20 neurons responding to two stimuli. In the first simulated scenario (Fig. 2A), only correlations between activity of different neurons, but not the single-neuron activities, are stimulus-modulated and thus encode stimulus information. The single cell information is zero, but the pairwise population information is not. Positive co-information arises because of large synergy with negligible redundancy. The Information Breakdown reveals that all the synergistic information is due to stimulus-dependent correlations. Population decoding with SVM of the N=20 neurons reveals that large-population information can be accessed exclusively with a non-linear decoder, and that shuffling correlations destroys all information, confirming it is exclusively encoded by correlations.

**Figure 2:**
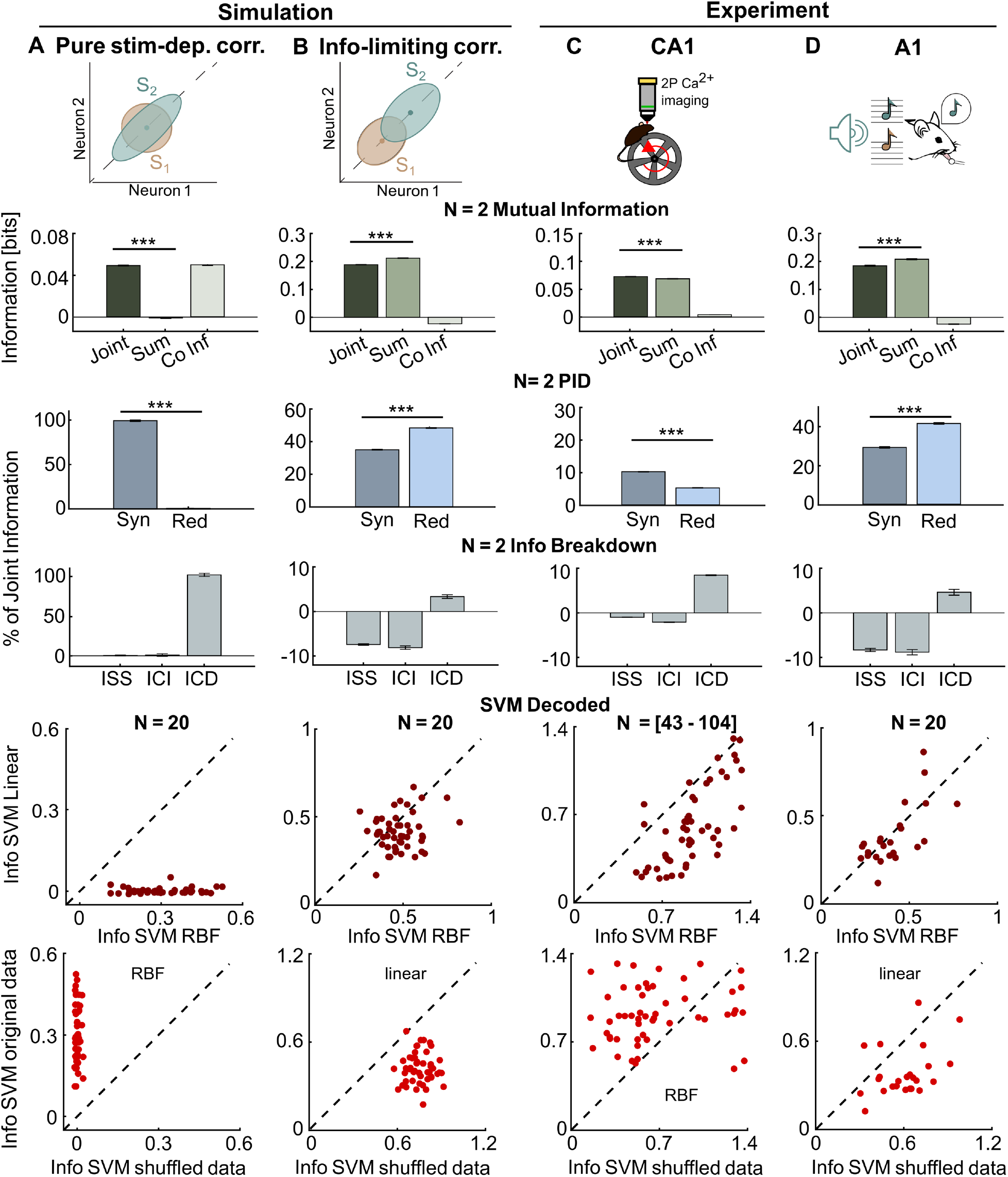
Assessing the role of correlations among neurons in neural population encoding. In each column we consider analysis of a different dataset. **A:** simulated population of N=20 neurons which carry information only by stimulus-dependent correlations, with no stimulus information provided by single-neuron firing rate modulation. **B:** simulated population of N=20 neurons which carry information by single neuron firing modulations and which have information-reducing correlations. **C:** CA1 recordings of N=[43-104] neurons over n=11 sessions during spatial navigation of a linear track in virtual reality. **D:** A1 recordings of N=20 neurons over n=12 sessions during tone presentation. In each row, we plot from top to bottom: direct calculation of information for neuron pairs and of sum of single neuron information; direct calculation of co-information for neuron pairs; direct calculation of synergy and redundancy for neuron pairs; direct calculation of the Information Breakdown components for neuron pairs; calculation of encoded information of the whole population using the information in the confusion matrix of an SVM decoder (linear or RBF), computed either on the real population responses (which contain correlations between neurons) or pseudo-population “shuffled” response obtained collecting randomly permuted trials to the same stimulus (shuffling removes correlations at fixed stimulus). In columns A-B we compute Shannon Information between neural activity and the identity of the two simulated stimuli. In column C-D we compute Shannon Information between neural activity and the identity of the presented tone (S=2 different tones) or the spatial location of the mouse (binning locations into S=12 equi-distant spatial bins), respectively. In column C, direct measures of pairwise information were obtained with R=2 equi-populated bins (appropriate for this dataset consisting of non-deconvolved calcium fluorescent traces). In column D, direct measures of pairwise information were obtained with R=3 bins, done by capping to 2 spike counts (appropriate for this dataset consisting of calcium signals deconvolved to estimate firing rates and activity counted in short windows). In each panel we plot mean ± SEM (for simulated data in panel A-B: over n=190 neural pairs and n=10 simulation repetitions for the direct information calculations; over n=5 different data folds and n=10 simulation repeats for the decoding information values; for CA1 data in Panel C: over n=10750 simultaneously recorded neuron pairs for the direct information calculations, and over n=11 recording sessions and n=5 trial folds for the decoding information values; for A1 data in Panel D: over n=2280 simultaneously recorded neuron pairs for the direct information calculations, and over n=12 recording sessions and n=2 trial folds for the decoding information values). Symbols *, **, *** denote two-tailed p<0.05, p<0.01, p<0.001 respectively, computed with paired t-tests.

In the second simulated scenario (Fig. 2B), information is encoded by single cells, correlations are only weakly stimulus-modulated, all neurons have equal stimulus tuning (responding more strongly to stimulus 2), and noise correlations are positive. In this configuration, redundancy is created (because all neurons have the same trial-averaged response profiles to the stimuli) and correlations reduce information (because they are elongated along the axis separating the mean firing rates of individual neurons and thus increase the overlap between the stimulus-specific distributions of neural activity) [20]. Negative co-information arises because of larger redundancy (created by so called signal similarity expressing the similarity of tuning to stimuli of individual neurons) than synergy (created by the small but present stimulus-modulation of correlations). Information Breakdown analysis reveals that indeed information is more redundant because the signal-noise similarity (captured by *I*_*sig*−*sim*_ and *I*_*cor*−*ind*_) is larger than the small stimulus-dependent correlations *I*_*cor*−*dep*_. In the large (N=20) population most information can be accessed with a linear SVM, with the non-linear SVM adding relatively little, and noise correlations reduce information (shuffling them away increases information).

We then applied the same analyses to two real neural datasets. We first analyze encoding of the mouse position (within a linear track) by populations of 43-104 simultaneously recorded neurons from the CA1 region of the mouse hippocampus (Fig. 2C). With the pairwise analysis, PID shows that both synergy and redundancy are present but synergy is larger and the Information Breakdown shows that this is due to modulation of the noise correlations strength with the position (*I*_*cor*−*dep*_∼10% of the pairwise information). Using a nonlinear decoder of the whole population increases information by ∼80% over what could be achieved with linear decoders, and shuffling data to destroy correlations decreases the nonlinearly decoded information by ∼80%, revealing a large effect of hippocampal noise correlations in position encoding by large neural populations, whose size could not be inferred by neuron pairs analysis.

We then analyzed encoding of sound intensity by populations of 20 neurons simultaneously recorded from the mouse auditory cortex A1 during pure-tone sound presentation (Fig. 2D). These networks were selected, among all recorded neurons, based on their encoding of task-relevant information in [90]. With the pairwise analysis, PID shows that both synergy and redundancy are present, but redundancy is larger. Information Breakdown analysis shows that this is due to negative *I*_*sig*−*sim*_ (neuron pairs have similar tuning to the stimuli) and *I*_*cor*−*ind*_ (most neural pairs have also positive correlations), with *I*_*cor*−*dep*_ contributing much less. Decoding whole-population activity with a nonlinear SVM did not increase the information decoded with a linear SVM (stimulus-dependent correlations were weak), and shuffling away noise correlations increases information substantially (thus correlations strongly reduced information).

Together, these results illustrate the power of combining MINT tools to understand deeply how interaction between neurons shape neural population coding.

### Computing how stimulus information encoded in neural population activity informs behavioral discriminations

Traditional approaches to neural information encoding of sensory stimuli have focused solely, as in the above examples, on how neurons or populations encode information about these stimuli. However, it could be that little or none of the information they encoded is actually utilized to inform behavior. It is thus important to have instruments to understand how much information in neural activity contributes to behavior.

Intersection Information (II) measures how much of the sensory information encoded in neural population activity is read out to inform behavior (Fig. 3A), and is computed with PID (using the tri-variate probabilities of stimuli, neural activity and behavioral choices) as the component of neural information that is both about stimulus and choice [24, 42, 43]. To demonstrate its use, we applied it to analyze the activity of populations of neurons recorded with 2-photon calcium imaging in mice in auditory cortex (A1) during pure-tone perceptual discrimination (Fig. 3B).

**Figure 3:**
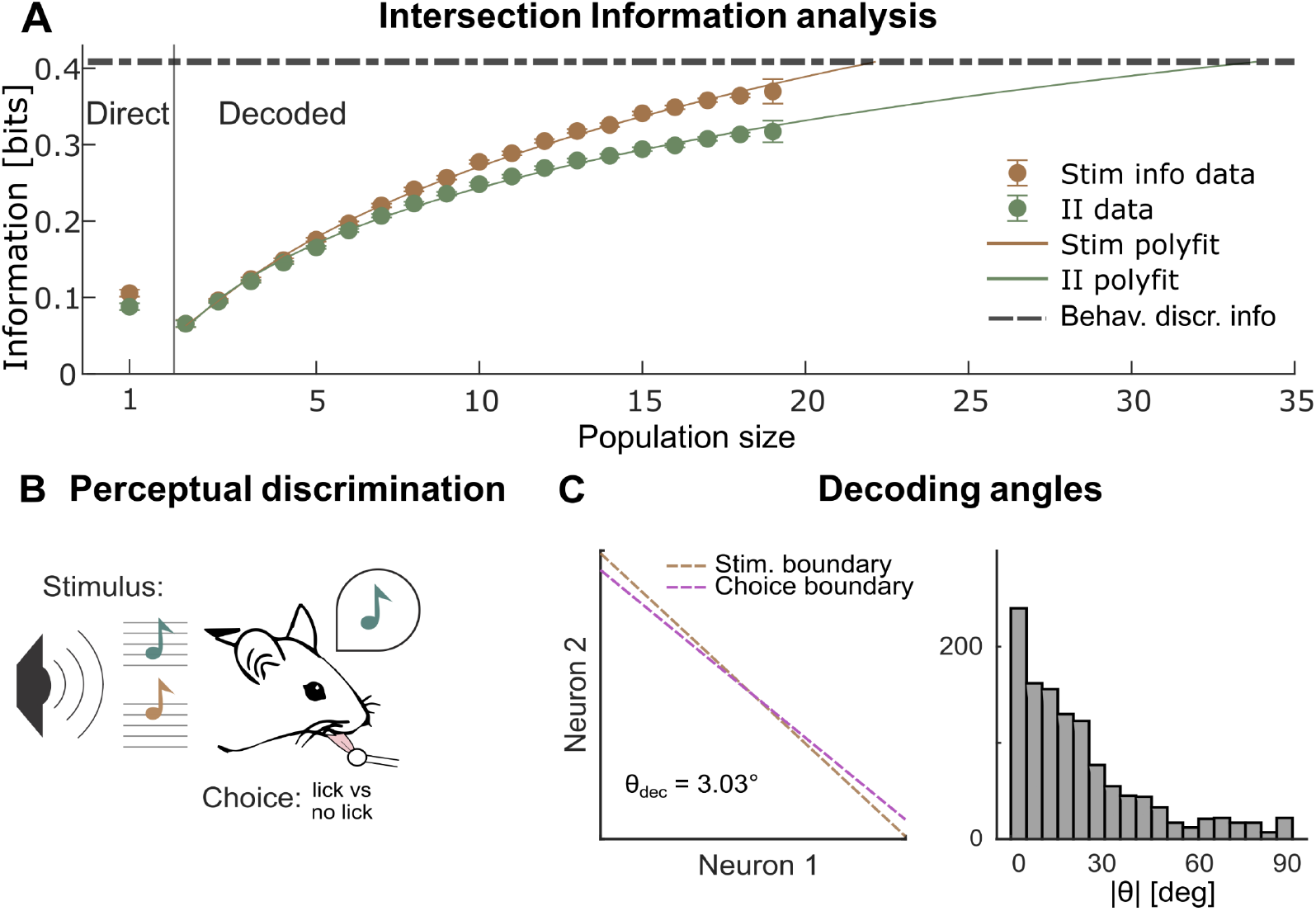
Stimulus and Intersection information coding in populations of cortical auditory neurons during a tone discrimination task. **A:** Stimulus information and Intersection Information encoded in neural activity recorded during a sound tone discrimination task. Left: single cell estimates using the direct method. Right: estimates of the information quantities using a RBF SVM (2-fold cross validation) as function of the population size. We plot the mean and SEM over all n=12 Field of Views and over all folds and over all subpopulations used. For population sizes N=2-18, for >100 independent subpopulations can be obtained, we shortened computation time using only n=100 randomly sampled subpopulations. For population size N=1 and 19, we used all the n=20 different subpopulations available. For the direct information calculation, we used 3 bins for 0 spikes, 1 spike and any value above 1. For all information analyses, we used the shuffle subtraction to correct for the limited-sampling bias. The dashed horizontal line plots the averaged information needed to explain behavioral discrimination accuracy (computed as the information between stimulus and choice). Full lines show log-polynomial fits to the dependence of stimulus and intersection information on population size. The population size with information sufficient to explain behavioral discrimination accuracy is the x-axis intercept of the point at which the fit lines cross. **B:** Schematic of the behavioral task in mice used when recording the data analyzed in this figure. **C:** Stimulus and choice boundary computed with MINT in the space of paired neural activity for one example neural pair in the dataset. The value of the angle between the two axes is reported in the inset. Right: distribution of the absolute value of the angle between the stimulus and choice boundaries for the n=2280 neural pairs in this dataset. See Supplemental Information SM7.2 and SM7.3 for details of simulations and real data analysis.

We first considered information encoded in single neurons, computed with the direct method. If the readout of the stimulus information in neural activity was optimal (respectively, completely suboptimal), II would equal the stimulus information, (respectively, be zero). We found that for single neurons, II was ∼90% of the total single-neuron stimulus information, showing that information encoded by these neurons is not read out optimally but still efficiently.

For sampling reasons explained above, the direct calculation of II can be done for small (N=1-3), but not for large populations. How can we use II to address how information relevant to behavior scales with population size? Specifically, how large must a population be to account for perceptual discrimination ability? To answer this, in MINT we combined II with dimensionality-reduction techniques. In this application, we used an SVM to compress neural activity (using `svm_wrapper.m` before `II.m`). This compression loses some information (the information values obtained with the direct method are ∼20% higher than the single cell values obtained with SVM decoders; Fig. 3A). However, II population information computed with SVM decoders are scalable and data-robust (Fig. S3). Computing how information scales with population size (Fig. 3A) shows that as population size increases, the gap between stimulus information and II widened. This means that behaviorally-relevant information is more redundant across neurons compared to information that is not used to inform behavior, confirming the usefulness of redundancy for behavioral readout [91]. Had we considered only stimulus information, we would have incorrectly concluded that ∼23 such neurons are sufficient to account for the mouse discrimination performance (Fig. 3A). However, taking intersection information into account reveals that ∼34 such neurons are instead needed to fully account for the perceptual discrimination ability, as not all stimulus information encoded in neural populations is read out (Fig. 3A).

We endowed MINT with instruments to characterize neural mechanisms of readout. Suboptimality may arise because of a misalignment between how information is encoded and how the brain reads it out to inform choices [42]. MINT returns the axes in neural activity space trained to discriminate between stimuli and the axes trained to discriminate different choices (using `svm_wrapper.m,` see Fig. S4 for examples on simulated data). Computing decoding angles of pairs of A1 neurons (Fig. 3C) shows that most pairs had a small but non-zero mis-alignment between stimulus and choice decoders, which explains the efficient but sub-optimal readout.

In sum, combining intersection information and dimensionality reduction can give precise insights about the behavioral relevance of information encoded by neural populations.

### Mapping content-specific encoding and transmission of information within a network

MINT provides both algorithms to study information encoding in individual network nodes and information transmission across nodes. We here illustrate how to combine them for reverse-engineering the information flow within neural networks.

We first simulated a network with four nodes *X*_1_, …, *X*_4_ each modeling the aggregate activity of a brain area (as e.g. measured by aggregate neural signals such as LFPs, M/EEG or fMRI). This network has a well-defined ground-truth flow of information about two independent stimulus features *S*_1_, *S*_2_ (Fig. 4A). Information about *S*_1_ is received from the outside by nodes *X*_1_and *X*_4_, and is then sent from *X*_1_ to *X*_2_ and *X*_3_. Information about *S*_2_ is received from the outside by *X*_2_which then sends it to *X*_1_. *X*_3_ and *X*_4_ exchange information which is not about *S*_1_ or *S*_2_. To disentangle the information flow, we computed (using the direct method) information encoded or transmitted at each time (in Fig. 4 we plot for each node and link the maximal information values over time, but we show in Fig. S5 that time-resolved analysis reconstructs correctly the ground-truth communication timing and delays), and we used MINT’s non-parametric permutations tests to identify significant encoding or transmission. Using Mutual Information between individual stimulus features and individual node activity reveals correctly that all nodes have information about *S*_1_ and that only *X*_1_ and *X*_2_ have information about *S*_2_ (Fig. 4B). To study how this information is exchanged within the network, we first computed overall information transfer with Transfer Entropy, finding correctly significant transfer from *X*_1_ to *X*_2_ and *X*_3_, from *X*_2_ to *X*_1_, and from *X*_3_ to *X*_4_ (Fig. 4C). To reveal the information content of this exchange we computed Feature Specific Information Transfer (FIT), revealing correctly that the information transferred from *X*_1_ is about *S*_1_ but not about *S*_2_, and that the information transferred from *X*_2_ is about *S*_2_ (Fig. 4D). FIT finds no information transfer from *X*_3_ to *X*_4_ about *S*_1_ or *S*_2_, thus determining correctly that the overall information transfer from *X*_3_ to *X*_4_ detected with TE is not about any of the two stimulus features. Finally, the finding that *X*_1_ and *X*_4_ encode information about *S*_1_ while they do not receive it from other network nodes implies that *X*_1_ and *X*_4_ receive external *S*_1_ information. Similarly, because *X*_2_ encodes information about *S*_2_ while not receiving within-network *S*_2_ information demonstrates that *X*_2_ receive external *S*_2_ information. Thus, combining encoding with transmission analyses could correctly reverse engineer the within-network specific information flow.

**Figure 4.**
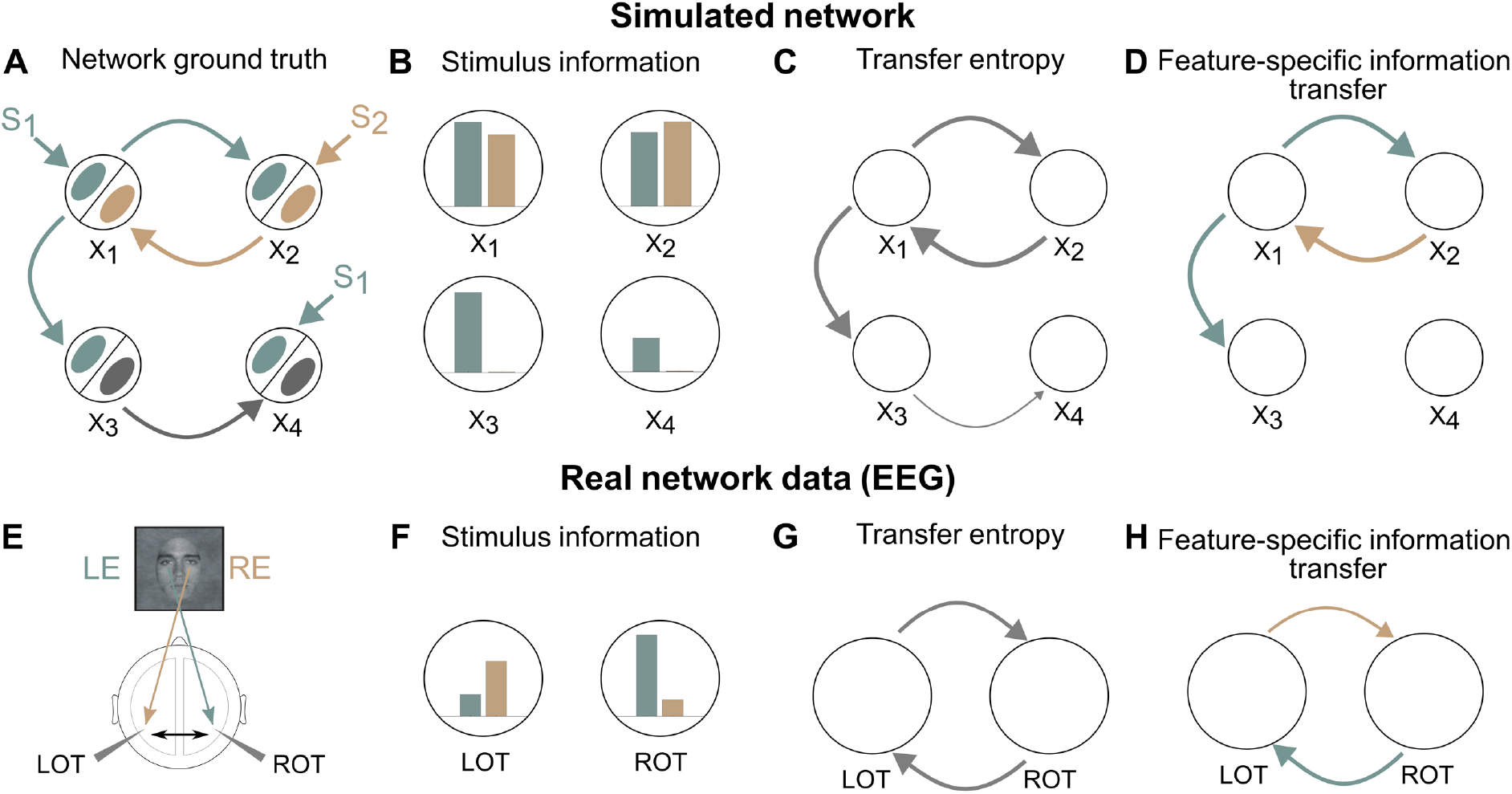
Reverse engineering information flow using stimulus-encoding and stimulus-transfer estimation algorithms. Panels A-D test MINT on simulated network data. **A:** Schematic of the simulation. The network comprises four neural nodes (black circles) *X*_1_,.., *X*_4_, each containing two subpopulations (ellipses within the circles) encoding two independent binary stimulus features *S*_1_, *S*_2_. The ground-truth stimulus specific information communication is plotted in Panel A, with grey color used to indicate no stimulus selectivity, and green and brown colors used to indicate information selectivity to *S*_1_ and *S*_2_ respectively. **B:** Maximum Mutual Information across time between each neural population *X*_*i*_ and the stimuli *S*_1_ and *S*_2_ **C:** Transfer Entropy (TE) between nodes. **D:** FIT about *S*_1_ and *S*_2_ between nodes. In panels C-D, only significant (p<0.01, permutation test) links are plotted, with thickness proportional to the computed value. In each panel we plot the average information values across n=10 simulation repeats. Panels E-F test MINT on real human EEG data. **E:** Schematic of the putative information flow inter-hemispheric information flow. LOT (ROT) denote Left (Right) occipito-temporal regions. LE (respectively RE) denote the Left (respectively Right) Eye face visibility feature. **F:** Maximum Mutual Information across time about the left or right eye visibility present in left of right OT region. **G:** Significant Transfer Entropy between LOT and ROT brain regions. **H:** Significant FIT between LOT and ROT brain regions. In panels G-H, only significant (p<0.01, permutation test) links are plotted, with thickness proportional to the computed value. In each panel we plot the average information values across n=15 experimental subjects. See Supplemental Information SM6.3 and SM7.1 for details of simulations and real data analysis.

We next tested how MINT reverse-engineers information flow in real brain networks by applying it to an existing EEG dataset recorded from human participants detecting the presence of either a face or a random texture from images covered by random bubble masks [92]. Prior work [36, 92, 93] revealed that the visibility of the eye region (proportion of visible pixels in the eye area) is critical for successful face discrimination and that the Occipito-Temporal (OT) EEG electrodes are those encoding most Mutual Information about both left and right eye visibility (Fig. 4E,F). To understand if some of this information was exchanged across the OT regions in different hemispheres, we used TE and FIT to analyze transmission of left or right eye visibility information across OTs. TE across hemispheres was found in both directions (right-to-left and left-right), suggesting a bi-directional inter-hemispheric communication (Fig. 4G). However, specific information transfer was precisely directional: FIT about the left eye was only from right-to-left and FIT about the right eye was only from left-to-right (Fig. 4H). Thus, using MINT allowed establishing encoding and directional transfer of different eye features across hemispheres with high specificity. These analyses could also temporally localize both encoding and inter-hemispheric transfer (Fig. S6).

Together, these results illustrate the power of combining MINT tools to reverse-engineer encoding and flow of specific information across brain networks.

### Availability and Future Directions

MINT is downloadable in source code (https://github.com/panzerilab/MINT with DOI 10.5281/zenodo.13998526), including a Dockerfile, and is licensed under GNU GPLv3. It contains documentation on using it and on building and installing it from source, unit tests, use examples, and replication of paper figures (https://github.com/panzerilab/MINT_figures).

The modularity of MINT allows it to be used alongside any other MATLAB function or toolbox. As exemplified above, we already provide pipelines for interfacing with decoding toolboxes. We plan to add plugins to generate neural and behavioral data from data acquisition and preprocessing toolboxes (e.g. [94]) with MINT’s input-data format requirements, and to generate MINT’s outputs suitable to be fed directly into toolboxes for further advanced analyses, e.g. for network analysis of information-transfer outputs [95].

MINT’s current version focuses on using discretized maximum likelihood estimators. We plan to endow them with optimal discretization algorithms based on model selection techniques (Akaike and Bayesian information criterions) as well as to endow them with other probability estimators including Maximum Entropy estimators [61], binless and nearest-neighbor estimators [96-98], and parametric probability models (Gaussian, Poisson) proposed in the neuroscience literature. The derivation of new neuroscience-related information quantities with PID is highly active [69, 99] and the open source and modularity of MINT will allow rapid integration of new developments. We are also developing a translated python version of MINT to widen usage.

## Acknowledgements

We thank C. Becchio, G. Iurilli and H. Nili for feedback. This work was supported by the Simons Foundation for Autism Research Initiative (SFARI) grant 982347 (to SP) and the NextGenerationEU grant FAIR PE0000013 (to TF). The Funders had no role in study design, data collection and analysis, decision to publish, or preparation of the manuscript.

## Data availability Statement

All data and code used to produce this paper is made publicly available. All code is downloadable in source code (https://github.com/panzerilab/MINT with DOI 10.5281/zenodo.13998526) and is licensed under GNU GPLv3. The neural CA1 data are attached to this submission as Supplemental Material.

## Supplementary Material

### SM1 Comparison with other toolboxes

The following table provides a synthetic comparison of main features of different currently available toolboxes.

**Table 1:** Comparison of MINT with other existing information-theoretic toolboxes. Abbreviations: FIT=Feature-specific information transfer; II= Intersection Information; MI= Mutual Information; PID=Partial Information Decomposition; TE = Transfer Entropy.

| Software | Information Measures | Input Data | Signif. Testing | Estimation Methods | Bias corr | Dim Red | Platform |
| --- | --- | --- | --- | --- | --- | --- | --- |
| MINT (Lorenz et al., this paper, 2024) | Entropy, MI, TE, Info Breakdown, PID, II, FIT Cond. variants | Discrete Continuous | Non-parametric permutation tests | Binning, Gaussian fit | ✓ | ✓ | MATLAB |
| FRITES (Combrisson et al., 2022) | Entropy, MI, TE | Discrete Continuous | Non-parametric, group stats | Binning, Gaussian copula | ✓ | X | Python |
| infoTheory (Climer and Dombeck, 2021) | MI (SMGM estimator) | Discrete Continuous | X | X | X | X | MATLAB |
| Infotheory (Candadai, 2019) | Entropy, MI, PID | Discrete Continuous | X | Binning Kernel-based | X | X | Python C++ |
| BROJA-2PID (Makkeh et al., 2018) | PID | Discrete | X | X | X | X | Python |
| Neuroscience Information Theory Toolbox (Timme and Lapish, 2018) | Entropy, MI, TE, PID, Info transfer, Cond. variants | Discrete Continuous | X | Binning | X | X | MATLAB |
| dit (James et al., 2018) | Entropy, MI, PID | Discrete | X | X | X | X | Python |
| Gaussian Copula Mutual Information (Ince et al., 2017) | Entropy, MI | Discrete Continuous | X | Gaussian copula | ✓ | X | MATLAB Python |
| Inform (Moore, 2017) | Entropy, MI, TE | Discrete | X | Binning | X | X | C (Python, R, Julia, Mathematica wrappers) |
| JIDT (Lizier, 2014) | Entropy, MI, TE, Cond. variants | Discrete Continuous | Non-parametric | Binning, Kernel-based Gaussian fit | ✓ | X | Java (Python, MATLAB wrappers) |
| MuTE (Montalto et al., 2014) | TE | Continuous | Non-parametric | Binning Gaussian fit Kernel-based | ✓ | X | MATLAB |
| ITE Toolbox (Szabó, 2014) | Entropy, MI | Discrete Continuous | X | Kernel-based | X | X | MATLAB Python |
| Trentool (Lindner et al., 2011) | TE | Continuous | Non-parametric, group stats | Kernel-based | ✓ | X | MATLAB |
| Transfer Entropy Toolbox (Ito et al., 2011) | TE | Spike trains | X | X | X | X | MATLAB |
| Information Breakdown Toolbox (Magri et al. 2009) | Entropy, MI, TE Info Breakdown | Discrete Continuous | Non-parametric | Binning, Gaussian fit | ✓ | X | MATLAB |
| PyEntropy (Ince et al., 2009) | Entropy, MI, Max entropy | Discrete Continuous | X | Binning, Shrink estimator | ✓ | X | Python |
| STAToolkit (Goldberg et al., 2009) | Entropy, MI | Spike trains | Non-parametric | Binning | ✓ | X | MATLAB (with .mex files) |

### SM2 Description of installation and testing of MINT, and of information theoretic tools implemented in MINT

MINT can be downloaded at the public repository https://github.com/panzerilab/MINT.

*Documentation on building and installing the software from source* are provided as a README file that specifies the installation requirements, as well as a build file (`buildMINT.m`) designed to automate the software’s compilation process. *Instructions on how a user can test the software on supplied simulated test data* are provided as a folder “How_to_use_MINT” in MINT’s public repository containing detailed instructions for testing it on simulated data. We also provide an additional repository (https://github.com/panzerilab/MINT_figures) containing the code that replicates all analyses in all figures, on both real neural data and simulated data. The dataset with CA1 neural data is provided as attachment in Supplemental Material, and the dataset with A1 neural data is provided by the public link https://drum.lib.umd.edu/items/30d43732-7149-4726-a860-0ae3d210b2ae.

MINT provides information theoretic tools to give quantitative answers to questions about information processing when applied to single neurons, population of neurons, or to aggregate neural signals recorded across multiple areas (including LFPs, M/EEG, fMRI). The considered information processed by the considered neural activity can be about a specific task variable, such as a sensory stimulus, a behavioral output, or about the activity of other neurons or neural populations.

All the information theoretic quantities are functions of the joint probability, sampled across experimental trials, of observing a given value for a set of task variables (e.g. sensory stimuli, movement parameters, behavioral choices) ***s*** ∈ ***S*** and of neural responses (***r***_**1**_, …, ***r***_***N***_) ∈ {***R***_**1**,…,_***R***_***N***_}. Each of the variables is indicated with bold font because it can be a multidimensional vector. Importantly, each dimension in the task and neural response variables is assumed to have discrete values, so that the probabilities can be estimated by empirical occurrences. In many cases, neural data will be already discrete in nature (for example, spike counts) and the same applies to some categories of task variables like behavioral choices or identity of the presented stimulus (which experimentally usually fall into a number of discrete categories). In other cases, either task variables or neural responses will be continuous data (e.g. LFPs, etc). These input data will be automatically discretized by MINT to perform the information calculations by specifying discretized into a finite number of bins by defining the number of bins `n_bins` (by default, 3 bins) and the binning strategy `bin_method` (including equi-spaced binning, equi-populated binning, and binning with user-defined bin edges; by default, no binning) as fields within the options input structure `opts`. MINT allows the use of these discretization procedures for all its information theoretic measures, and direct probability distribution sampling after discretization was used for all results in the paper. In addition to binning, MINT offers the option to compress the multi-dimensional neural activity space (***r***_**1**_, …, ***r***_***N***_) ∈ {***R***_**1**,…,_***R***_***N***_} into a dimensionality reduced representation, obtained with either supervised decoding methods or unsupervised data reduction methods, which can be also discretized and used for information calculations with the direct probability distribution sampling (see Section SM5, Fig. S2).

The functions to compute the information quantities in MINT follow a consistent structure (Fig. S1). The first input provides the data organized in a cell array. This input should be formatted as {A, B, C, …}. Optionally, the second input can be a cell array of strings called `reqOutputs`, which specifies the quantities to be computed based on the provided datasets, following the nomenclature of the input format. As a last input the user can provide a structure `opts` that contains optional arguments for the computation.

The outputs of the functions are also implemented in a consistent structure. The first output variable contains cells with the requested information quantities given in `reqOutputs` (in the same order as specified). The second output variable contains cells with the naive information quantities (i.e., no limited-sampling bias correction) and the last output variable contains the null distribution for each specified information quantity, if the `opts` field `computeNulldist` is set to true. For instance, to compute the limited-sampling bias corrected and the naive Mutual Information between two populations X1 and X2 (two-dimensional arrays with neurons in the first dimension and trials in the second dimension) the MI function is called as follows:

`[MI_corr, MI_naive] = MI({X1, X2},{‘I(A;B)’},opts)`

If the input data for `H.m`, `MI.m`, `cMI.m`, `PID.m` or `II.m` is given as a time series (three dimensional array, with neuron or brain area ID in the first dimension, time points in the second dimension, and trials in the third dimension), these functions compute the information quantities for each time point and output them as time series.

In the following we list and synthetically describe the information quantities implemented in MINT.

#### SM2.1 Shannon information

The `MI.m` function computes Shannon Mutual Information between a population of *N* neurons {***R***_**1**_, …, ***R***_***N***_} and a task variable ***S*** (such as a sensory stimulus). It is a non-parametric measure that quantifies the full single-trial relationship between {***R***_**1**_, …, ***R***_***N***_} and ***S***. It is defined as [3]:

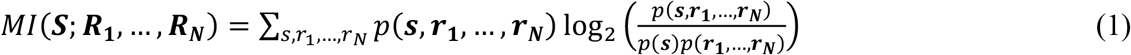

where *p*(***s, r***_**1**_, …, ***r***_***N***_) is the joint probability, sampled across experimental trials, of observing stimulus value *s* ∈ *S* and the neural responses (***r***_**1**_, …, ***r***_***N***_) ∈ {***R***_**1**_, …, ***R***_***N***_}, and *p*(***s***) and *p*(***r***_**1**_, …, ***r***_***N***_) are the marginal probabilities of observing *s* and (***r***_**1**_, …, ***r***_***N***_), respectively. The sum in Eq. (1) spans all possible events. *MI*(***S***; ***R***_**1**_, …, ***R***_***N***_) is non-negative and is zero if and only if {***R***_**1**_, …, ***R***_***N***_} and ***S*** are independent. To compute the Shannon Mutual Information of two variables, `reqOutputs` has to be defined as `‘I(A;B)’`. Moreover, MINT also allows to compute the mutual information between ***S*** and {***R***_**1**_, …, ***R***_***N***_} conditioned on the activity of another population of M neurons {***R***′_**1**_, …, ***R***′_***M***_} or another stimulus feature ***S***′ (function `cMI.m`).

#### SM2.2 Information Breakdown

The `MI.m` function can also compute measures that quantify how interactions between neurons contribute to the encoding of ***S***. The desired quantities can be passed to the function as specific strings within the `reqOutputs` cell array input. These quantities include the co-information ***coI*** (`reqOutputs` defined as **‘**coI(A;B)**’**), which is defined as the difference between the population information and the sum of single-neuron stimulus information quantifying the overall contribution of interactions to population encoding. Additionally, the function can return Information Breakdown terms [21] quantifying how pairwise correlations contribute to co-information. These terms include the signal-similarity *I*_*sig*−*sim*_ (contribution of the similarity across neurons of trial-averaged responses to different stimuli, `reqOutputs` defined as **‘**Iss(A)’), the stimulus-independent correlations *I*_*cor*−*ind*_ (contribution of the interplay between the signs of signal similarity and of noise correlations, `reqOutputs` defined as **‘**Ici(A;B)**’**), and the stimulus-dependent correlations *I*_*cor*−*dep*_ (quantifying how much information is gained by the stimulus-modulation of noise correlations, `reqOutputs` defined as **‘**Icd(A;B)**’**).

#### SM2.3 Partial Information Decomposition (PID)

The `PID.m` function computes PID components. PID decomposes the information jointly carried by a set of source variables (for us, a set of simultaneously recorded neurons; the first N variables in the `inputs` cell array) about a target *S* (for us, a sensory stimulus or a behavioral choice; the last variable in the `inputs` cell array) into non-negative components that capture information about the target that is either redundantly encoded across sources, uniquely encoded by a single source or synergistically encoded by the combination of sources. For more than two source variables such components can also represent combinations of redundancy, synergy and unique information. In the case of two source variables, once the individual Mutual Information between the target and each source and the joint Mutual Information between the targeted and the two sources, and one of the PID components (e.g. redundancy) are all computed, algebraic linear relationships (derived from PID “lattices”) allow to compute all remaining PID components. Thus, a PID with two sources is defined by the choice of a specific redundancy measure. The desired PID components can be passed to the function as strings within the `reqOutputs` cell array input (`‘Red’` for redundancy, `‘Syn’` for synergy, `‘Unq1’` and `‘Unq2’` for the unique information carried by the first or the second input source, respectively). For more than two variables, defining a redundancy measure is enough to compute each PID component as a linear combination between redundancy terms (the output name `‘PID_atoms’` provides all PID components). Therefore, different methods to decompose information differ and are defined by the measure of redundancy they use (which can be specified as the `redundancy_measure` field in the `opts` structure). In MINT we implemented three possible measures of PID redundancy, which are very popular and respect the so called pairwise marginal property (redundancy is invariant for distributions preserving the pairwise marginals between each source and the target).

##### *SM2.3.1 I*_min_

We implemented Williams and Beer’s PID original redundancy measure called *I*_*min*_. This measure [68] quantifies the information redundantly encoded about ***S*** across N source variables {***R***_**1**,…,_***R***_***N***_} as:

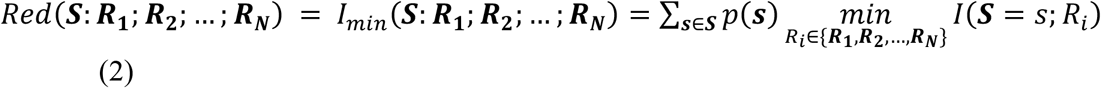

where *I*(***S*** = ***s***; ***R***_***i***_) represents the specific information that source variable ***R***_***i***_ provides about a specific value ***s*** of the target variable ***S***. *I*_*min*_ captures redundancy as the similarity across the source variables ***R***_***i***_ in distinguishing individual values of ***S***.

##### SM2.3.2 Minimum Mutual Information (I_MMI_)

MINT also implements the PID based on the redundancy measure introduced in [71], called *I*_*MMI*_. This measure quantifies the information redundantly encoded about ***S*** across N source variables {***R***_**1**,…,_***R***_***N***_} as:

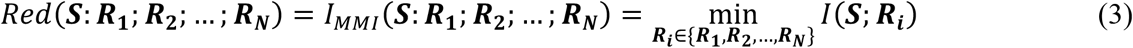

Therefore *I*_*MMI*_ captures redundancy as the minimum amount of information encoded about ***S*** by any of the source variables. When applied to Gaussian variables, its results coincide with the ones calculated through the *I*_*min*_ (SM2.3.1) and the BROJA definitions (described below, SM2.3.3).

##### SM2.3.3 BROJA

Finally, MINT implements the PID based on the redundancy measure termed BROJA [70], which defines redundancy about ***S*** between two source variables ***R***_**1**_ and ***R***_**2**_ as the result of a constrained optimization problem:

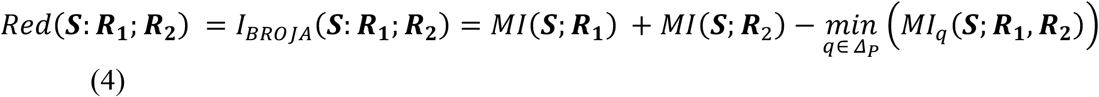

where the minimum of the Mutual Information is taken over the space *Δ*_*P*_ of distributions preserving pairwise marginals between individual source variables and the target variable. BROJA’s advantage is that it is additive for independent systems of sources and targets, however it is only defined for two source variables. The numerical calculation of Eq. (4) is performed by MINT though the conic optimization Embedded Conic Solver (ECOS) algorithm [52]. The use of ECOS requires the installation of a C library which can be implemented by either compiling the C source code with a MATLAB-compatible C Compiler (using MINT’s `BuildMINT.m` function) or copying locally the precompiled binaries that MINT provides for Linux, Windows and macOS.

#### SM2.4 Information transmission measures

MINT provides a number of measures to quantify the overall or feature-specific directed information transmitted from a putative sender region ***X*** to a receiver region ***Y*** from which neural activity was simultaneously recorded. These measures are all based on the Wiener-Granger causality principle which states that a region ***X*** causally influences a region ***Y*** if the past state of ***X*** predicts the present state of ***Y*** at time t beyond what can be predicted by the past state of ***Y***.

##### SM 2.4.1 Transfer Entropy (TE)

Transfer Entropy from ***X*** to ***Y***, *TE*(***X*** → ***Y***) [72], measures the overall information transmitted from region ***X*** to region ***Y*** and can be computed using the `TE.m` function in the toolbox. *TE*(***X*** → ***Y***) is defined as the conditional Mutual Information between the past activity of the sender region ***X***_*past*_ and the present activity of the receiver region ***Y***_*pres*_ given the past of the receiver ***Y***_*past*_. Moreover, MINT also allows to compute TE from ***X*** to ***Y*** conditioned on the activity of a third node ***Z* (**function `cTE.m`; ***Z*** can, in principle, also be the multivariate activity of a set of regions).

Although in the example computations of TE provided in Figs 4, S5 and S6 we computed the present of *Y* at a single time point t and the past of ***X*** and of ***Y*** at individual time points lagged by a delay Δt, MINT also allows computing information transfer measures for present and/or past activity as multidimensional variables, potentially spanning several time points.

The string to define in `reqOutputs` to compute Transfer Entropy from the data in the first to the data in the second element of the `input` cell is `‘TE(A->B)’`. In addition, a second output can be requested with `‘TE(B->A)’`, in order to reduce computational effort and exploring data more efficiently by calculating both directions of Transfer Entropy with a single function call.

##### SM 2.4.2 Feature-specific information transfer (FIT)

Feature specific information transfer from ***X*** to ***Y*** about a stimulus feature *S FIT*(***S*** → ***X*** → ***Y***) measures the information transmitted from ***X*** to ***Y*** about a specific feature ***S*** and can be computed using the `FIT.m` function in the toolbox. *FIT*(***S*** → ***X*** → ***Y***) is defined as the minimum between two PID terms with similar but slightly different interpretations. The first term is the information about ***S*** that is redundant between ***X***_*past*_ and ***Y***_*pres*_, and is unique to any information encoded by ***Y***_*pres*_. The second term is the information about the present activity of the receiver ***Y***_*pres*_ that is redundant between ***X***_*past*_ and ***S***, and its unique to any information encoded by ***Y***_*past*_. To guarantee the nonnegativity of FIT, both terms are computed using the *I*_min_ measure REF. Minimizing between the two terms ensures key FIT properties, including that it is upper bounded by the feature information encoded by the past of region *X I*(***S***; ***X***_*past*_), the one encoded by the present of region ***Y*** *I*(***S***; ***Y***_*pres*_) and by the overall information transmitted from ***X*** to ***Y*** *TE*(***X*** → ***Y***). Moreover, MINT allows for the computation of conditional FIT (cFIT, using the `cFIT.m` function), to remove from *FIT*(***S*** → ***X*** → ***Y***) the component potentially routed through the past activity of a third recorded region ***Z*** (***Z*** can, in principle, also be the multivariate activity of a set of regions). *cFIT*(***S*** → ***X*** → ***Y***|***Z***) is defined as *FIT*(***S*** → ***X*** → ***Y***) minus a term capturing feature information transmitted from ***X*** to ***Y*** that is also redundantly encoded by the past of ***Z*** [36].

Similar to the TE function, the FIT function can compute bidirectional information transfer about a target by specifying `‘FIT(A->B;S)’` and `‘FIT(B->A;S)’` in the `reqOutputs` cell array. The same applies to cFIT in `cFit.m` (specifying `‘cFIT(A->B;S|C)’` and `‘cFIT(B->A;S|C)’` in the `reqOutputs` cell array).

#### SM2.5 Intersection Information

MINT implements also Intersection Information (II, computed using the `II.m` function), a measure quantifying the amount of sensory information encoded by neural activity that is used to inform behavioral choices. II quantifies the part of information in neural responses that is common to both stimulus and choice information. II is computed as the minimum between two PID terms with similar but slightly different interpretations. The first term is the information about choice ***C*** redundant between stimulus ***S*** and neural response ***R***. The second term is the information about stimulus ***S*** redundant between choice ***C*** and neural response ***R***. By default, these terms are computed using the BROJA redundancy measure. Minimizing between the two terms ensures that II satisfies key properties that would be expected from a measure with this interpretation, including that independent ***S*** and ***R*** imply null intersection information, that II satisfies the data processing inequality, and that II is upper bounded by both *MI*(***R***; ***S***) and *MI*(***R***; ***C***).

To compute the Intersection Information as the amount of information present in input data A, encoded in input data B that is also present in input data C, `reqOutputs` can be defined as `‘II(A,B,C)’`.

### SM3 Limited-sampling bias correction

MINT implements several limited-sampling bias corrections of information theoretic quantities. For Shannon Entropy, Mutual Information and for the Information Breakdown terms, bias-corrected estimates are computed separately for each quantity. For the PID calculation, bias-corrected estimates can be obtained either by correcting each PID atom individually, or alternatively by correcting for the bias first the individual and joint mutual information term and one of the PID terms (e.g. redundancy or synergy), and then by correcting for the bias of the other PID components by using algebraic relationships derived from PID “lattices” (see SM 3.7). By default, no bias correction is computed (uncorrected, or naïve, information quantities).

#### SM3.1 Shuffle Limited-sampling bias correction

When this bias correction option is called (setting the `bias` field in the options structure `opts` to `‘shuffSub’`), the bias is estimated by computing the information values after destroying all genuine stimulus information by randomly permuting the stimulus-neural response association in each trial [80,81], and then it is subtracted out from the original estimate to provide the bias-corrected estimate. MINT allows to subtract these bias estimates from all implemented information theoretic quantities. A non-zero integer in the field `shuff` of the options structure `opts` specifies the number of shuffles to be performed and averaged over to obtain this estimate (by default, 20 shuffles).

#### SM3.2 QE correction

The Quadratic Extrapolation (QE) procedure [73,80] (setting the `bias` field in the options structure `opts` to `‘qe’`) assumes that the estimation is performed in a regime with large numbers of trials and approximates the bias of the information quantities as a second-order expansions in the inverse of the number of available trials [78]. This procedure first re-computes the information from fractions (halves and quarters) of the data available and then fits the dependence of information estimates on the inverse number of trials to a quadratic function. This quadratic fit is then used to estimate the bias-corrected information value as the value that would be obtained with the quadratic scaling law if an infinite number of trials were available (i.e., the intercept term of the fit). MINT allows the use of this bias correction with all its information theoretic measures. Field `xtrp` in the options structure `opts` specifies the number of repetitions of the extrapolation procedure. The function performs the specified number of extrapolations and calculates the final corrected value as the mean of the estimates (by default, 10). Note that MINT allows the option to perform a linear (rather than quadratic) extrapolation, which uses only halves and not quarters of the data. This may be convenient when very small datasets are available and the division into quarters is problematic.

#### SM3.3 Shuffle-QE correction

MINT also implements a bias correction combining both Shuffle and QE (setting the `bias` field in the options structure `opts` to `‘qe_shuffSub’`). This procedure first computes both the original and the shuffled values, performs QE on both as explained above and then computes the unbiased estimate by subtracting out the QE-corrected shuffled value from the QE-corrected non-shuffled value. The parameters `shuff` and `xtrp` can be set in the options `opts` structure as mentioned above.

#### SM3.4 Panzeri-Treves correction

This correction technique analytically approximates the linear term of the bias expansion in the inverse of the number of available trials, which is then subtracted from the measured information to obtain bias-corrected information values [78,80]. The estimation of the bias depends only on the number of response bins with a non-zero probability of being observed, which is estimated using a Bayes approach. It is available only to Shannon Entropy and Mutual Information (setting the `bias` field in the options structure `opts` to `‘pt’`) but not to PID-based quantities.

#### SM3.5 Ish bias reduction procedure for multi-dimensional data

The *I*_*sh*_ procedure is relevant for the reduction of the bias of the information about a task variable (say stimulus S) carried by the joint observation of a multivariate neural response with dimension N (e.g. the activity of N neurons). It adds and subtracts to the definition of Mutual Information two entropy terms which have equal asymptotic value (in case of exact sampling of the probabilities with an asymptotically large number of trials). For limited number of trials, the difference between these two terms provides a negative contribution to the Mutual Information bias. Thus, *I*_*sh*_ has a considerably smaller bias (though larger variance) than the direct estimate of Mutual Information from Eq. (1). Another interesting property of *I*_*sh*_ is that it typically has a negative bias, whereas direct estimates of Mutual Information typically have a positive bias. Thus, the joint calculation of Mutual Information from Eq (1) and *I*_*sh*_ allows the estimation of upper and lower bounds to the real information values. This procedure can be applied to the Mutual Information and to the Information Breakdown (where it allows computation of upper and lower bounds of some terms) (setting the `bias` field in the options structure `opts` to `‘shuffCorr’`). It can be combined with the QE and the shuffle-subtraction corrections.

#### SM3.6 Best Universal Bound procedure

This method developed by [79] expresses the information estimation as a polynomial approximation problem, allowing the computation of ‘Best Universal Bounds’ on the information bias and variance (setting the `bias` field in the options structure `opts` to `‘bub’`). Its results depend on the selection of a parameter k_max_ related to degrees of freedom and specified in the field `k_max` in the options structure `opts(`by default, 10).

#### SM3.7 Options for PID bias

For QE, shuffle subtraction, and shuffle-QE subtraction applied to PID with two source variables, two bias-correction options are available. The first option is to correct for the bias, with the chosen bias correction procedure, each PID term individually. The second option is to correct for the bias in a way that respects the linear relationships between the PID terms and Shannon information quantities derived from the so called PID lattices. This is done by correcting for the bias the individual and joint mutual information term and one PID component of choice, and then correct for the bias the other PID components by using the algebraic relationships derived from PID “lattices”. The chosen PID atom can be specified with field `chosen_atom` in the options structure `opts(`by default, synergy). This option is implemented by default (setting the `opts` field `‘pidConstrained’` to true). If one chooses not to, each PID atom is corrected individually.

### SM4 Hierarchical permutations for testing statistical significance and the impact of correlations among different data dimensions in information coding

#### SM4.1 Data shuffling

Shuffling neural data is a useful tool to test hypothesis and gain insights in the information encoding structure of neural populations and how information is transferred. The `hShuffle.m` function provides a range of hierarchical data shuffling methods, allowing for disruption of neural correlations, temporal patterns or stimulus information. Each data feature (a neural activity dimension or a task variable) can be shuffled either unconditionally on any other variable or conditionally on the values of other variables. For example, neural responses can be shuffled unconditionally on any other variable to destroy the information they carry about the stimulus (e.g., to shuffle the first two input variables across trials the `reqOutputs` can be defined as `‘AB’` and the `opts` field `‘dim_shuffle’` is set to `‘Trials’`). Alternatively, neural responses of different neurons can be shuffled conditional on the stimulus values to provide surrogate data that preserve single-neuron stimulus information but that destroy noise correlations (correlations at fixed stimulus between different neurons). Shuffling the neural responses across trials with the same stimulus but keeping the position of each timepoint fixed provides surrogate data that keep time-resolved stimulus information of single neurons but disrupt across-time correlations. For example, to shuffle the first input variable conditioned on the second and third input variables the `reqOutputs` cell can be defined as`’A_BC’` (the variable(s) to condition the shuffling on are specified after the underscore) and the `opts` field `‘dim_shuffle’` is set to `‘Trials’`.

Shuffled data are either used within a given function (e.g. `MI.m` or `FIT.m`) to output null hypothesis values, or to be separately provided as input to any function in MINT to construct user-defined non-parametric null distribution to empirically estimate the p-value of measures computed from the original data.

Furthermore, MINT enables efficient computation of group-level averaged shuffled quantities by recombining shuffled information values across experiment repetitions using the `create_nullDist_groupLevel` function. This function accepts measures calculated from M independent data shuffles across N experiment repetitions and outputs K distinct realizations of the permuted group average.

#### SM4.2 Cluster permutation for multiple comparison correction

It is often important to detect significant information encoded or transmitted across data points that are correlated due to physical proximity (e.g., in space and time, or time and communication delay). MINT implements the rigorous detection of clusters of adjacent significant information values via cluster permutation tests [48] (`clusterStatistics.m` function). The `clusterStatistics.m` function takes as input a matrix of information values computed from the original data across adjacent space and time points and a set of M analogous matrices obtained from shuffled data. The function computes a cluster-forming threshold as a specified percentile `clusterPercentilThreshold` (provided as input) of the shuffled information values. This threshold is calculated either by pooling shuffled values across all samples (when the input `pool` parameter is set to 1) or independently for each sample (when `pool` is set to 0). The latter option provides less statistical power but is recommended when shuffled information is non-stationary over space and time. The procedure then identifies clusters in both the original and shuffled data by connecting adjacent information values that surpass the cluster-forming threshold and computes the mass of each cluster as the sum of its information values. A cluster-level null distribution is created by taking the maximum cluster mass from each shuffled dataset. Finally, clusters in the original dataset that exceed a specified percentile `significanceThreshold` (provided as input) of this null distribution are classed as significant.

### SM5 Interfacing information calculations with dimensionality reduction methods

To allow information analysis on datasets with a high number of dimensions, we offer several possibles wrappers that integrate tools or dimensionality reduction in a way that permits the calculation of information measures on processed variables with less dimensionality.

#### SM5.1 Interfacing information calculation with supervised dimensionality-reduction method

Here we describe which supervised dimensionality reduction algorithm we implemented in MINT and we interfaced then with the information theoretic algorithms.

Our routines take input data in the form of a cell array with neural responses (***r***_**1**_, …, ***r***_***N***_) ∈ {***R***_**1**,…,_***R***_***N***_} across all trials in the first element and the task variables ***s*** ∈ ***S*** (e.g. sensory stimuli, movement parameters, behavioral choices) across all trials in the second element. It returns an array across all trials of dimensionality reduced representation 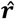 of (***r***_**1**_, …, ***r***_***N***_). The dimensionality-reduced representation 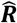 is computed by cross-validated decoding of the behavioral variables ***s*** given the neural data, so that the representation 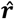 for each trial is computed from trials held out from the decoder’s training process. The reduced neural representation data are then fed as neural data input to the information calculation routines.

The reduced representation 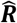 obtained through the supervised decoding method can be understood as a representation of the neural activity data which is lower dimensional (typically, one dimensional) but that still captures efficiently the information about ***S*** provided by the joint neural responses. The reduced representation 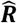 can take form of the value of the behavioral variable decoded as most likely from neural activity (which can be directly fed to the information calculation routines, an approach that is equivalent to computing information from the confusion matrix of the decoder), or the posterior probability of the task variable value given the considered neural activity (which can then be binned and fed to the information calculation routines).

Mutual Information between the stimulus and the reduced neural representation 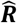, and Intersection information between stimulus, choices, and the reduced neural representation 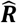 are obtained by simply using the reduced neural representation 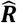 instead of the actual neural activity ***R*** into the corresponding equations defined in sections SM 2.1 and SM2.5. Importantly, both the Mutual and the Intersection Information satisfy the data processing inequality, which implies that the values obtained through the reduced neural representation are lower bounds to the values of the full, unreduced, neural activity representation.

Supervised models implemented in MINT are:

##### SM5.1.1 Support Vector Machines (SVM)

Support vector machines (SVM) are supervised machine learning methods that find the optimal hyperplane/s in the data space (in our case neural data (***r***_**1**_, …, ***r***_***N***_)) to classify the labels (in our case ***s***). MINT provides a function `svm_wrapper.m` that trains and tests the SVM with either linear or radial basis function (RBF) kernel, using either `fitcsvm.m` (for binary labels; Statistics and Machine Learning Toolbox in MATLAB), `fitcecoc.m` (for multiclass labels; Statistics and Machine Learning Toolbox in MATLAB) or the libsvm toolbox [84] as the underlying SVM implementation. The first element of the output cell array is the lower dimensional representation 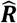 as an array of the cross-validated predicted labels across trials. The second element of the output cell array is the representation 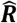 as an array of cross-validated posterior probabilities of ***s*** given the neural responses in that trial. The function can also output the weights of the decoding model for the linear SVM (which can be used e.g. to calculate the angle between boundaries as in Fig. S4) or the trained hyperparameters, in case they were optimized.

##### SM5.1.2 Generalized linear model (GLM)

Generalized linear models are an extension of linear models that can incorporate non-Gaussian-distributed data (including discrete data). In MINT, the `glm_wrapper.m` function trains and test GLM models using lasso, ridge or elastic net regularization and either `lassoglm.m` (Statistics and Machine Learning Toolbox in MATLAB) or the `glmnet` toolbox [85]. This function allows for the training and testing of GLM models with optional regularization methods, including lasso, ridge, or elastic net. It provides flexibility in choosing regularization types and additional options for k-fold cross-validation. Depending on the option chosen in `reqOutputs`, it outputs as 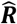 an array of cross-validated predicted labels for each trial, or cross-validated posterior probabilities of ***s*** given the neural data in each trial.

#### SM5.2 Interfacing information calculation with unsupervised dimensionality-reduction method

Unsupervised methods transform neural data into a lower dimensional output 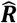 that still approximate the data well. Our routines take input data in the form of a list across all trials of neural responses (***r***_**1**_, …, ***r***_***N***_) ∈ {***R***_**1**,…,_***R***_***N***_}, as well as the desired dimensionality of the reduced representation. It returns a list of reduced representation 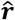 of (***r***_**1**_, …, ***r***_***N***_). The reduced neural representation data are then fed as neural data input to the information routines for information calculation. Unsupervised models implemented in MINT are describe below.

##### SM5.2.1 Principal Component Analysis (PCA)

Principal Component Analysis (PCA) is a dimensionality reduction technique that uses an orthogonal linear transformation to project the data onto a lower dimensional space with maximal variance (REF). MINT’s function `pca_wrapper.m` outputs in each trial the coefficients of the data along the selected number of principal components.

##### SM5.2.2 Non-negative Matrix Factorization (NMF)

Non-negative Matrix Factorization (NMF) is another technique of dimensionality reduction that projects the data onto a lower-dimensional space that still describes the data well. Unlike PCA, it does not require different components to be orthogonal, but requires that the decomposition is performed with nonnegative coefficients and basis functions (which is recommended for reducing the dimensionality of inherently nonnegative data, such as spike counts). MINT’s function `nmf_wrapper.m` outputs in each trial the non-negative coefficients of the data along the selected number of principal components.

### SM6 Details of simulations

#### SM6.1 Simulations of neural populations information encoding

This section presents a detailed description of the simulation and the analysis of information encoding in neural populations presented in Fig. 2.

We simulated two scenarios which capture two main ways in which correlations have been reported to influence population coding [1,21,87]. For each scenario we simulated correlated spike trains of a neural population of N = 20 neurons responding to two simulated stimuli (200 trials per stimulus, 10 simulation repetitions).

The strength of correlations between neurons was modulated by generating responses to each stimulus as the sum of an independent Poisson process (independent outcome for each neuron) and a shared Poisson process (same outcome across neurons), adjusting the pairwise Pearson noise correlation for each stimulus by varying the contribution of the shared and independent processes to the spike trains. Thus, the spike count of neuron *i* (*i* = 1,2) was generated as:

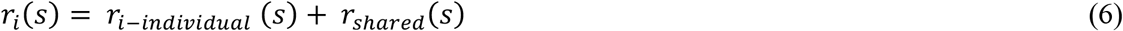

where *r*_*i-individual*_ (*s*) and *r*_*shared*_(*s*) are the output of 3 independent Poisson processes for each stimulus, with mean count parameter indicated by the corresponding name.

The first scenario (Fig. 2A) was implemented with strong stimulus modulation of the correlation strength, resulting in information-enhancing noise correlations. The individual and shared processes were created such that the resulting total firing rate of each of the neurons is constant across stimulus values, so only the firing correlation is informative about the stimulus. The parameters were *r*_*i-individual*_ (*s* = 1) = 1 *sp*/*s, r*_*shared*_ (*s* = 1) = 1 *sp*/*s, r*_*i-individual*_ (*s* = 2) = 2 *sp*/*s, r*_*shared*_ (*s* = 2) = 0 *sp*/*s*.

For the second scenario (Fig. 2B) we simulated information limiting noise correlation. Namely we simulated a population of neurons all with the same stimulus selectivity (lower spiking rate to the first stimulus and a higher spiking rate to the second stimulus and thus positive signal correlations) and with positive noise correlations that were only weakly stimulus dependent. The parameters were *r*_*i-individual*_ (*s* = 1) = 0.8 *sp*/*s, r*_*shared*_ (*s* = 1) = 0.2 *sp*/*s, r*_*i-individual*_ (*s* = 2) = 1.9 *sp*/*s, r*_*shared*_ (*s* = 2) = 0.1 *sp*/*s*.

The so generated spikes counts were binned into 5 bins, by leaving spike counts ≤ 4 untouched and setting to 4 all spike count values ≥ 5 (this was done by setting input options `opts` of `MI.m` the `binning_method` field `‘userEdges’,` that allows binning the data with user-defined bin edges). All MI and PID values were corrected for the limited-sampling bias by using the shuffle-subtraction procedure implemented in the toolbox (averaged over 30 shuffles).

We used the `svm_wrapper.m` function of the toolbox to predict the stimulus based on the population activity by fitting a cross-validated Support Vector Machine (SVM) with 5 folds. Two distinct kernel functions were employed to fit the SVM to the data: linear and RBF. We performed hyperparameter optimization, tuning the parameters C for linear SVM and C and gamma for SMV RBF using 2 folds cross-validation. We used Bayesian optimization over a logarithmic scale ranging from 10^−3^ to 10^3^ and maximum number of iterations equals 30. To evaluate the role of correlations in information encoding we computed the Mutual Information of the predicted and the true stimuli. To eliminate noise correlations, pseudo-responses were generated by shuffling the simulated response conditionally on the stimulus value, so they have the same single cell properties as the original data but no noise correlations (h`Shuffle.m` function, `reqOutputs` defined as`’A_B’` and `‘dim_shuffle’` set to `‘Trials’`). We computed the Mutual Information using the `MI.m` function of the toolbox between the actual stimulus and the one predicted from all above-described decoders (linear and RBF kernel SVM for simulated response and pseudo-response) and compared them to gain insights into the effects of noise correlation on the population information. The simulation was repeated n = 10 times and results were averaged across repeats.

#### SM6.2 Simulation of encoding and readout of information in pairs of neurons

This section presents a detailed description of the simulation and analysis of information encoding and readout from pairs of neurons presented in Fig. S4.

We simulated a pair of neurons, independently encoding a binary stimulus *S* with values *s* ∈ [−1,1] across 1000 trials. The single-trial firing rate of each neuron *i* ∈ 1,2 was determined by a Poisson process *r*_i_(*S*)∼*Poisson*(*λ*_i_(*S*)), with the intensity parameter *λ*_i_(*S*) depending on the stimulus as *λ*_i_(*S*) = *λ*_0_ + Δ ⋅ *W*_*enc,i*_ ⋅ *S. λ*_0_ determined the mean firing rate of each neuron across trials, Δ the separation in the mean firing rates across the two stimuli, and *W*_*enc,i*_ was the element *i* of the 2-dimensional encoding vector 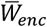 determining the tuning of each neuron to the stimulus. In our simulations we set *λ*_0_ = 4, Δ= 1 and 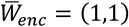, so that both neurons had lower firing rates for *s* = −1 and higher firing rates for *s* = 1.

On each trial, we simulated a choice variable *C* by taking the dot product between the population firing rates and a decoding vector 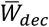, such that 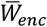 and binarized *C* using equi-populated binning. We obtained the decoding vector 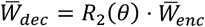 by applying the standard two-dimensional rotation matrix R_2_(*θ*) to the encoding vector 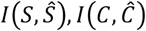 as follows 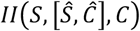.

We used the `svm_wrapper.m` function of the toolbox to train a cross-validated (5 folds) linear SVM (hyperparameter *C* = 1) to decode the stimulus *Ŝ* and the choice *Ŝ* from the neural population activity. We computed stimulus, choice and intersection information as *I*(*S, Ŝ*), *I*(*C, Ĉ*) and *II*(*S*, [*Ŝ, Ĉ*], *C*) respectively, using the `MI.m` and the `II.m` functions of the toolbox (with no bias correction).

We simulated two different scenarios, one with a small (*θ* = 20°) and one with a large (*θ* = 70°) angle between the encoding and the decoding vectors. We computed the angle between the decision boundaries of the SVM trained to decode the stimulus 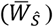 and the one to decode the choice 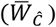 as

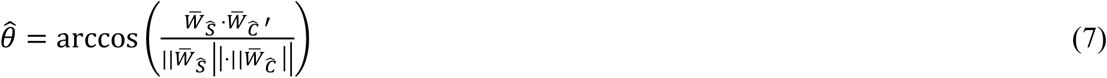

where the prime symbol (′) indicates the transpose operation, and 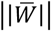 indicates the norm of vector 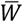. A total of 5 simulations were conducted for each scenario, information and angle values were averaged across simulations.

#### SM6.3 Simulation of aggregate activity signals in networks of interacting nodes

This section presents a detailed description of the simulation and analysis of aggregate signal activity in a network of interacting nodes (Fig. 4 and Fig. S5).

The network implemented content-specific encoding and transmission of information across four nodes, *X*_1_, *X*_2_, *X*_3_ and *X*_4_ (N = 4) each of them divided into two subnodes *X*_*N*,1_ and *X*_*N*,2_ (M = 2). A total of 10 simulations were conducted, each including 200 trials with a duration of 30 ms. We simulated a stimulus that included two binary features *S*_1_ and *S*_2_ whose values were drawn independently in each trial. The value of *S*_1_(*t*) was set equal to the value of *s*_1_ ∈ [−1,1] while the value to *S*_2_(*t*) was set equal to the value of *s*_2_ ∈ [−1,1] within a defined stimulus-active time window. Outside this window, the value was set to zero.

Subnodes *X*_1,1_ and *X*_4,1_ received input regarding *S*_1_, while subnode *X*_2,2_ received input regarding *S*_2_. The stimulus-active time window for *X*_1,1_ and *X*_2,2_ was defined as [3, 12] ms, while *X*_4,1_ received the input with a delay Δt_2_ of 12 ms (stimulus-active time window [15, 24] ms). In addition, all subnodes received stimulus-feature unrelated activity at any timepoint as zero-mean Gaussian noise *ε*_*N,M*_(*t*) = *N*(0, *σ*_*noise*_) with standard deviation *σ*_*noise*_ = 0.5.

To simulate the transfer of feature-related and unrelated activity from one subnode to another, a delay Δt_1_ of 5 ms was defined. With that time delay, subnode *X*_1,1_ transmitted its activity to *X*_2,1_ and *X*_3,1_, subnode *X*_2,2_ to *X*_1,2_ and *X*_3,2_ transmitted its activity to *X*_4,2_.

The aggregated activity of each node is defined as the summed activity of the two subnodes. The activity of the four nodes at each time point is defined as:

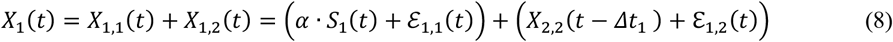

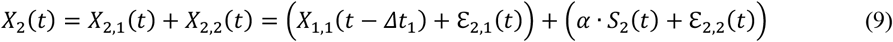

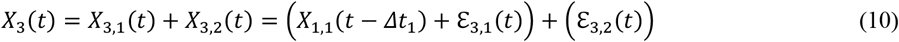

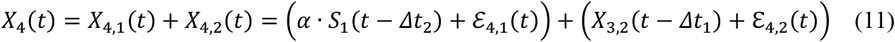

For all subsequent analysis we binned the activity of each node into R = 3 equi-populated bins and we corrected for the limited-sampling bias of information with the QE procedure implemented in the toolbox.

First, we computed the Mutual Information between each node and the stimulus features at all time points (see Fig. S5), using the `MI.m` function of the toolbox.

Consistent with the implemented stimulus input and information transfer, *X*_1_ exhibited information about *S*_1_ from 3 to 12 ms, while *X*_2_ and *X*_3_ displayed a 5 ms delayed stimulus-feature informative window. The simulated activity of *X*_4_ contained information regarding *S*_1_ from 15 to 24 ms, consistent with the implemented input delay of 12 ms compared to *X*_1_.

Information about *S*_2_ was only present in *X*_1_ ([8, 15] ms) and *X*_2_ ([3, 12] ms), in line with the implemented input of *S*_2_ and delayed transfer from *X*_2_ to *X*_1_.

To gain insight into the information transfer within the network we computed the transfer entropy between all pairs of simulated nodes, using the `TE.m` function provided by the toolbox. Based on the established ground truth, the temporal parameters to compute transfer entropy were defined as t = 12 ms and *Δt* = 5 ms. Consistent with the implemented network interactions, we found significant transmission of information from *X*_1_ to *X*_2_ and *X*_3_, from *X*_2_ to *X*_1_ and from *X*_3_ to *X*_4_ (see heatmap in Fig. S5A).

In the context of real data analysis, the optimal temporal parameters are typically not known. Therefore, we demonstrate in the next step how one can assess these parameters by computing the time-delay maps of information transmission between nodes *X*_1_ and *X*_2_. To compute the content of information flow, we measured FIT with the `FIT.m` function implemented in the toolbox. To obtain the temporal profile of content-specific information transmission and to reconstruct the delay of information transmission, we first computed FIT at each time step of the simulation with all possible delays for *X*_1_ → *X*_2_ related to *S*_1_, *X*_1_ → *X*_2_ related to *S*_1_ and for *X*_2_ → *X*_1_ related to *S*_2_ (see Fig. S5B). Consistent with the implemented ground truth, significant transfer of information was found with a delay of 5 ms from *X*_1_ to *X*_2_ related to *S*_1_ and from *X*_2_ to *X*_1_ related to *S*_2_.

By computing FIT for all pairs of nodes (t = 12 ms, Δt = 5 ms), we found significant information transfer related to *S*_1_ from *X*_1_ to *X*_2_ and *X*_3_ and significant information transfer related to *S*_2_ from *X*_2_ to *X*_1_ (see Fig. 4D and Fig. S5C).

To test for significance in the information theoretic quantities averaged across the n = 10 simulations, we used non-parametric permutation tests. For each simulation, we first conducted two different shuffling procedures 100 times and recomputed TE and FIT from the shuffled data (*t* = 12 ms, Δt = 5 ms). First, we conditionally shuffled the sender activity at fixed value of the stimulus to preserve stimulus induced covariations between the sender and the receiver and destroy single-trial correlations contributed by real communication. Second, we shuffled *S* for FIT to break any relationship between the stimulus and variables *X* and *Y*, and we shuffled *X* for the TE analysis to break any relationship between *X* and *Y*. We then took the pairwise maximum between the information values obtained from the two shuffling procedures to obtain a single, conservative null distribution [36]. Using the `create_nullDist_groupLevel.m` function of the toolbox we generated 500 samples of the null distribution of the permuted average across simulations, and estimated TE and FIT p-values empirically. To estimate the significance of FIT in the time-delay domain (Fig. S5C), we implemented the same procedure outlined above computing FIT at each time step of the simulation with all possible delays. We then used cluster permutation (with the `pool` option set to 0) setting both `clusterPercentilThreshold` and `significanceThreshold` to the 99^th^ percentile to individuate significant FIT clusters.

### SM7 Supplemental details and supplemental results of analyses of real neural data

#### SM7.1 EEG analysis methods

We analyzed a publicly available EEG dataset [92] (available at https://datadryad.org/stash/dataset/doi:10.5061/dryad.8m2g3). Full details are reported in the original publication. Here we summarize them briefly. The EEG data were recorded while participants (n=16) performed a face detection task. Participants were presented with an image hidden behind a bubble mask that was randomly generated in each trial. The presented image was an image of a face in half of the trials and a random texture in the other half of the trials. Participants were instructed to report whether a face was present or not. In our analyses, we only considered trials where the face was correctly detected by the participants (approximately 1000 trials per subject). Following the recommendations of the original publications analyzing these data [92,93], we excluded one participant from the analysis. All analyses in our paper are based on the n=15 selected participants. EEGs were recorded by fitting participants with a Biosemi head cap comprising 128 EEG electrodes. EEG data were re-referenced offline to an average reference, band-pass filtered between 1 Hz and 30 Hz using a fourth order Butterworth filter, down-sampled to 500 Hz sampling rate and baseline corrected using the average activity between 300 ms pre-stimulus and stimulus presentation. ICA was performed to reduce blink and eye-movement artifacts (see [92,93]).

For the analyses of TE and FIT, we selected the EEG electrodes in the left and the right Occipito-Temporal regions that had the highest Mutual Information about the visibility of the contra-lateral eye, exactly as done in previous papers [93,36]. We computed the first derivatives of the EEG signal for both Occipito-Temporal sensors and used both their absolute values and first derivatives to compute the information quantities, for consistency with analyses performed in previous studies [93,36]. As stimulus feature for the computation of Mutual Information and FIT, we used the visibility of an eye (defined as the fraction of pixels within the eye region that were not hidden by the bubble mask). Both neural and stimulus features were discretized using 2 equi-populated bins. We computed the information quantities for all combinations of directionality of flow across hemispheres (left to right, right to left) and eye identity (left or right eye). As done in previous papers REF to compute a single TE and FIT value for each participant we selected a rectangular region in the time-delay domain centered around the contra-lateral FIT peaks (time ranging from 140 ms to 240 ms peri-stimulus presentation, delay ranging from 20 ms to 90 ms; same for both eyes, as they were significant in very similar time-delay regions). We computed the average over delays and then picked the maximum over time within this region. We used the same procedure described in SM6.3 to compute the significance of the across-participants averaged FIT and TE (Fig. 4G-H), generating 500 null samples from 10 shuffles within each participant.

Files that reproduce the analysis of these data are found in Supplemental Material, file MINTfigures.zip, subfolders ‘Figure4’.

#### SM7.2 Analysis of CA1 data

We reanalyzed a previously published dataset [27] in which the activity of several tens to a few hundreds of neurons was recorded simultaneously using in-vivo two-photon calcium imaging from CA1 neurons in head-fixed transgenic mice during virtual reality navigation of a linear track. This dataset is provided as Supplemental Information file ‘CA1_data.mat’.

We analyzed neurons recorded from *n*_*FOV*_ = 11 Fields of View (FOV) from *n*_*A*_ = 7 animals. For consistency with the previous study reporting the original data [27], the spatial position of the linear track was computed by binning the space along the track into *S* = 12 equi-populated bins. Also, the neural activity *r*_i_ of each neuron was quantified by binning the calcium traces into R = 2 equi-populated bins (only raw calcium traces and not deconvolved signals were available from [27]. For the PID analysis, we used all individual neurons present in the dataset leading to *n*_*pairs*_ = 36158 pairs of simultaneously recorded neurons used for the pairwise direct information analyses and to *n*_*sessions*_ = 11 sessions for the population vector analyses. The neural responses dimensionality was reduced using linear and nonlinear (RBF) SVM to predict the position categories using 5-fold cross-validation and hyperparameter optimization (2-fold cross-validation, Bayesian optimization over a logarithmic scale ranging from 10^−3^ to 10^3^ for C and gamma and maximum number of iterations equals 30) and later the predicted labels were used jointly to replace the full neural response.

Files that reproduce the analysis of these data, as well as the neural data themselves, are found in Supplemental Material, file MINTfigures.zip, subfolder ‘Figure2’.

#### SM7.3 Analysis of A1 data

We reanalyzed a previously published dataset [90] in which the activity of several neurons was recorded simultaneously using in vivo two photon calcium imaging from A1 L2/3 neurons in head-fixed transgenic mice during a pure-tone discrimination task. Data are publicly available at https://doi.org/10.13016/m2yt-mfxk.

The experimental task was structured as follows. After a pre-stimulus interval of 1 s, head-fixed mice were exposed to either a low-frequency (7 or 9.9 kHz) or a high-frequency (14 or 19.8 kHz) tone for a period of 1 s. Mice were trained to report their perception of the sound stimulus by their behavioral choice, which consisted of licking a waterspout in the post-stimulus interval (0.5–3 s from stimulus onset) after hearing a low-frequency tone (target tones) and holding still after hearing high-frequency tones (non-target tones). Two-photon calcium imaging was used to acquire the calcium fluorescence signals from individual A1 L2/3 neurons during the task with an imaging frame rate of 30 Hz. We pre-processed these data as follows to match the pe-processing used by the authors in the original publication. We smoothed the raw calcium fluorescence traces using a zero-phase (MATLAB `filtfilt.m` function) order-2 low-pass Butterworth filter (`butter.m` function in MATLAB) with normalized cutoff frequency of f/(fs/2), where f=2Hz is the low-pass cutoff frequency, and fs=30 Hz is the sampling frequency of the calcium imaging data. As in the original publication [90] the resulting traces were deconvolved with a first-order autoregressive model.

We analyzed neurons recorded from n= 12 Fields of View (FOV) from n = 12 animals. For consistency with the previously published work, we only considered the 20 individual neurons in each session with the shortest-latency intersection information peak, as described in [90]. This led to selecting n = 2280 pairs of simultaneously recorded neurons used for the pairwise direct information analysis and n=12 sessions for the population analyses. For the information analysis of these data, we identified for each neuron the imaging time frame within the trial of maximal intersection information exactly as in the original publication [90].

We then considered for each neuron a time frame of n = 10 imaging frames (corresponding to a window of 333 ms) around the peak intersection information time frame (we call this the peak time window for the neuron). Then we discretized activity for each neuron into R = 3 bins according to whether it was detected 0, 1 or > 1 spikes in the peak time window (this was done by setting in input options `opts` of `MI.m` the `binning_method` field `‘userEdges’,` that allows binning the data with user-defined bin edges). The stimulus set used for the stimulus encoding analysis was binary, dividing the presented sound tones into the low-and high-frequency categories. The choice set used for the intersection information analysis was also binary (lick vs no lick). For Figure 3, we used the `svm_wrapper.m` function of the toolbox to train a cross-validated (2 folds) RBF SVM (hyperparameter *C* = 1) to decode the stimulus *Ŝ* and the choice *Ĉ* from the neural population activity. We computed stimulus and intersection information as Mutual Information *I*(*S, Ŝ*) between presented and decoded stimulus and II as the Intersection Information *II*(*S*, [*Ŝ, Ĉ*], *C*) between the presented stimuli, the mouse choices and the stimulus and choice decoded from neural activity using the `MI.m` and the `II.m` functions of the toolbox. The bias was corrected by applying the shuffle subtraction procedure setting the `shuff` field of the `opts` structure to 30. Figure S2 illustrates the pipeline of dimensionality reduction and information measurement used to generate Fig. 3. The fitting was performed using a second-order polynomial on the logarithm of the population size.

Files that reproduce the analysis of these data are found in Supplemental Material, file MINTfigures.zip, subfolders ‘Figure2’ and ‘Figure3’.

## Supplementary Figures and Figure Captions

**Figure S1.**
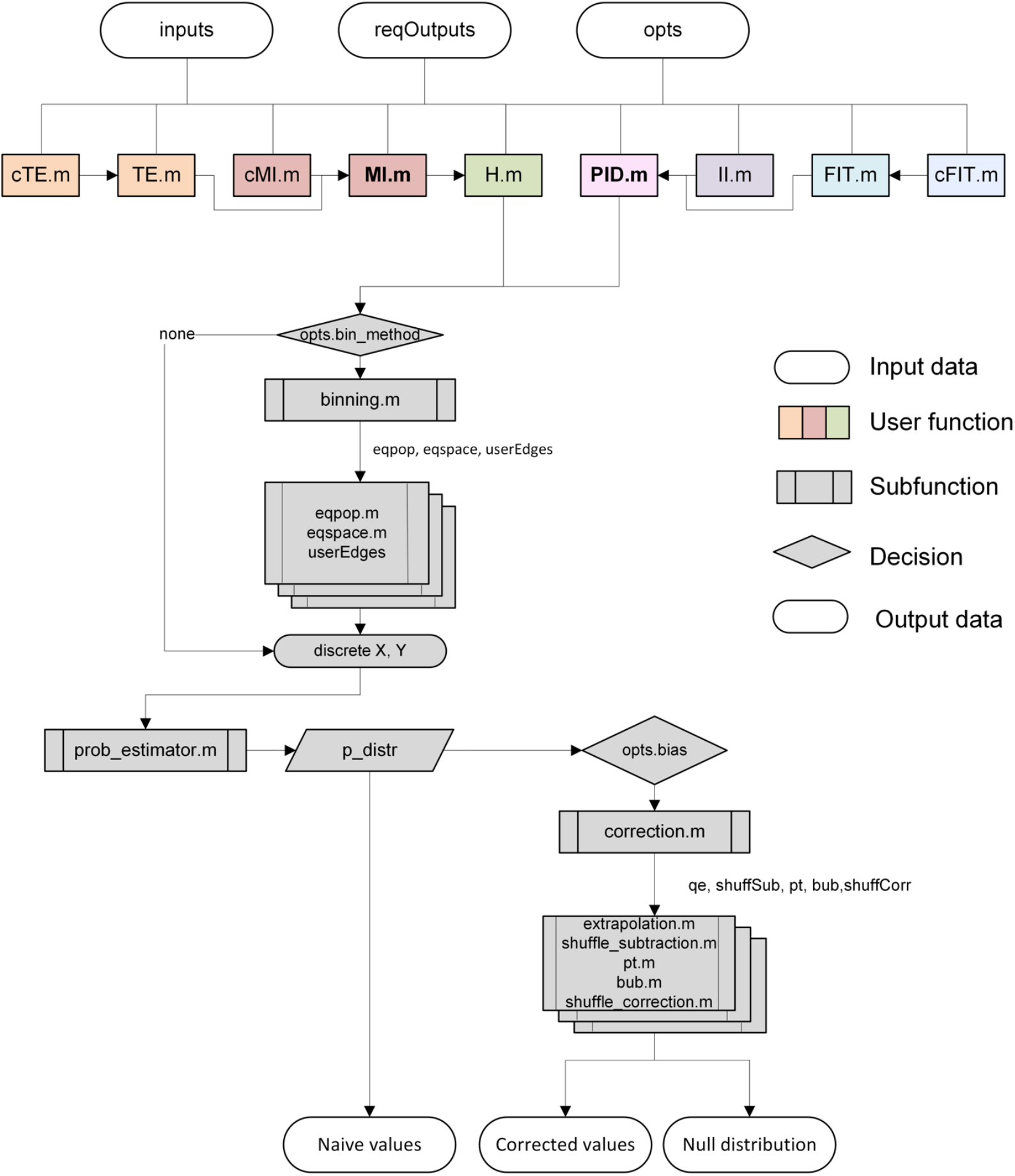
Flowchart of the MINT toolbox. The flowchart illustrates the structure and workflow of the MI module of the Toolbox, highlighting the steps involved in computing information values.

**Figure S2.**
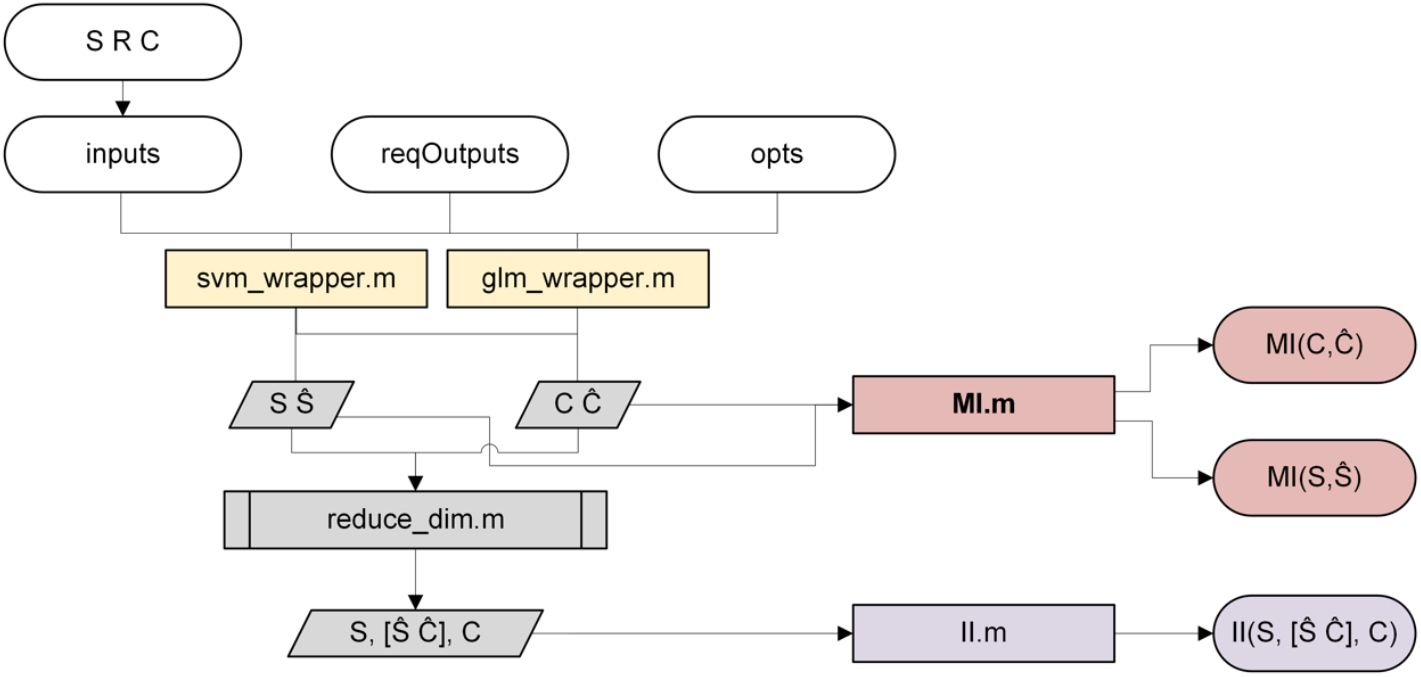
Example flowchart of the pipeline module used with the MI and II functions. This is an example pipeline using the dimensionality reduction wrappers in the toolbox with information-theoretic functions. One could also use the unsupervised wrappers (PCA or NMF) to do the dimensionality reduction or any other of the information functions (PID, TE, etc.).

**Figure S3:**
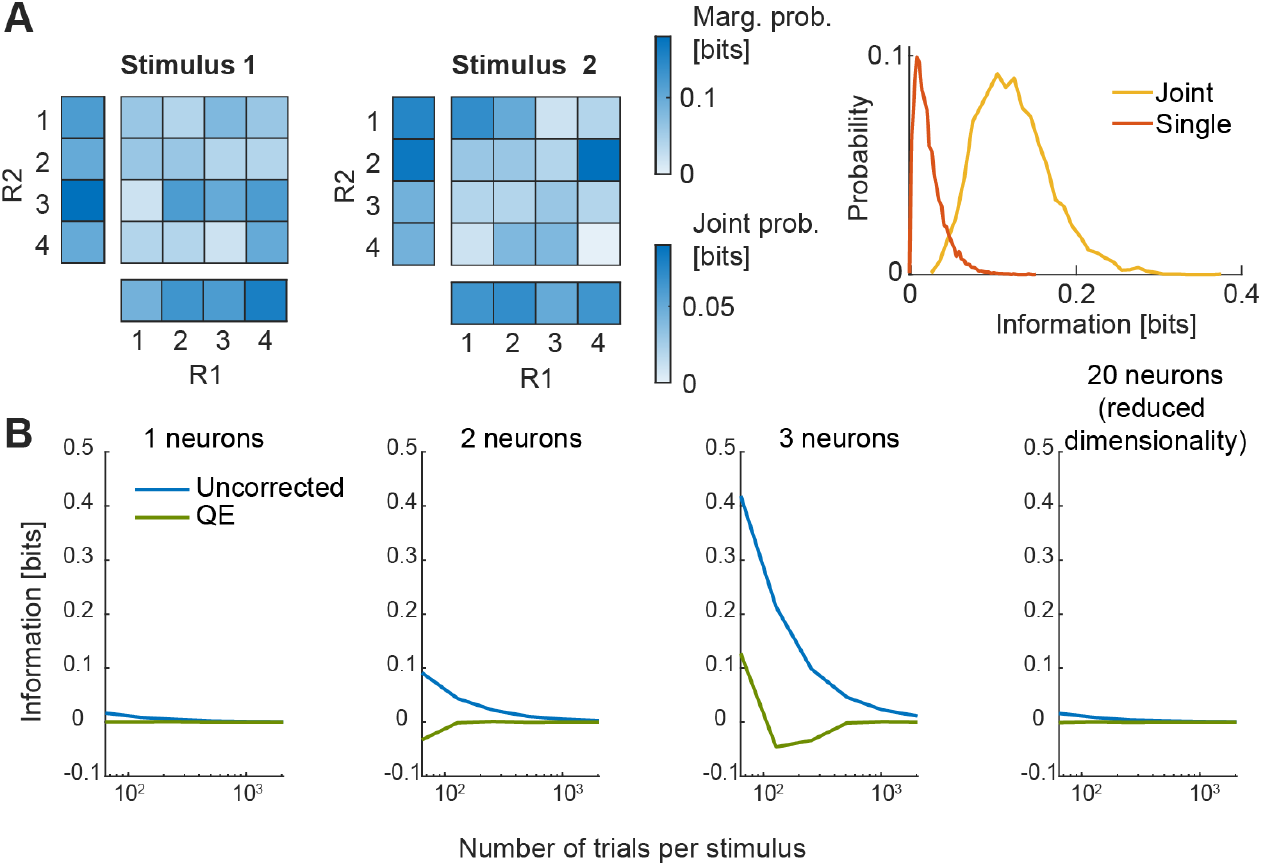
Limited-sampling bias of the direct method explodes with neural response dimensionality, but information can still be computed robustly with dimensionality reduction techniques. **A**. Schematic illustration of the limited-sampling bias problem. Two uninformative neurons, responding on each trial with a uniform distribution of spike counts ranging from 1 to 4, regardless of which 2 stimulus values were presented. The empirical response probability heatmaps sampled from 50 trials per stimulus are shown in the left and middle columns (responses to stimuli 1 and 2, respectively). Side heatmaps indicate the marginal probability values. Because of limited sampling, the neural response probabilities look different across stimuli (even if they are not), and much more so for the joint probability than for the marginal response probabilities. Right: distribution (over 5,000 simulations) of the plugin information values obtained with 50 trials per stimulus. Although the information should be zero, it is > 0 because of the random sampling variations illustrated in panel A, left. The bias is the (non-zero) average value of this distribution, as the true asymptotic value should be zero. The bias is larger for the joint information (when neural activity is 2-D) than for the single neuron information (when neural activity is 1-D). **B**. Average over 200 simulations, each performed with the number of trials per stimulus shown on the x axis, of values of Mutual Information *MI*(R; *S*) between a binary stimulus S and a neural response ***R*** made of *n* uninformative neurons. (Each neuron emits in each trial an integer value between 1 and 4 in a stimulus-independent uniform distribution like in panel A, left). Again, information here should be zero, so the value plotted corresponds to the limited-sampling bias, either without subtracting any bias correction (“uncorrected”) or after performing the QE bias correction. For 1, 2 and 3 neurons the information is calculated with the direct method, while for the 20 neurons it is first reduced to one dimension using the `svm_wrapper.m` cross-validated with 3 folds. The correction was done with 20 repetitions (`xtrp`=20). Using the direct method, the bias is under control with bias corrections with reasonable numbers of trials when considering one or two neurons, but it explodes out of control for 3 neurons. In contrast, the calculation of information from a population of 20 neurons can be performed robustly with limited data sizes when using dimensionality reduction.

**Figure S4:**
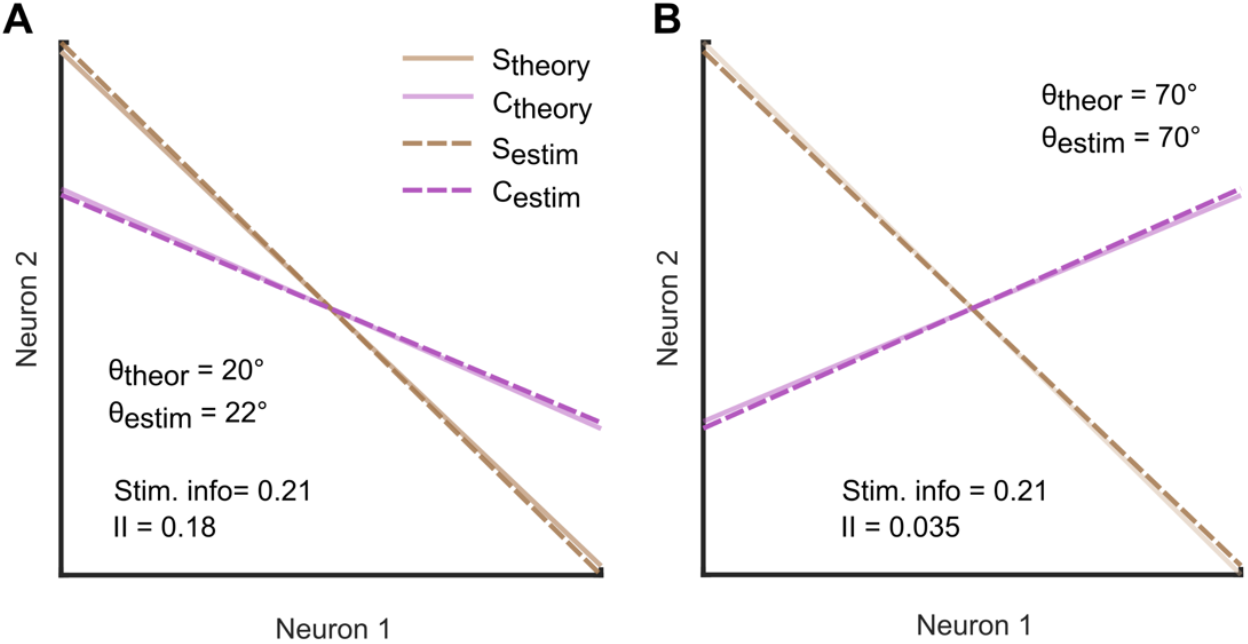
Role of stim-choice angle in the intersection information. Real and decoded stimulus and choice boundaries for a simulated population of 2 neurons with a small (A) and a large (B) angle between the stimulus and choice boundaries. The plot are made in the 2-D space of the firing rate of the two simulated neurons. This plot shows that MINT can reconstruct precisely the true neural activity axes that generate choices and stimulus coding. The figure also illustrates that, while stimulus information is the same for the two scenarios, the intersection information (II) is much larger for the case when stimulus and choice boundaries are well aligned [42]. Simulations performed as detailed in Supplemental Material Section SM6.2

**Figure S5.**
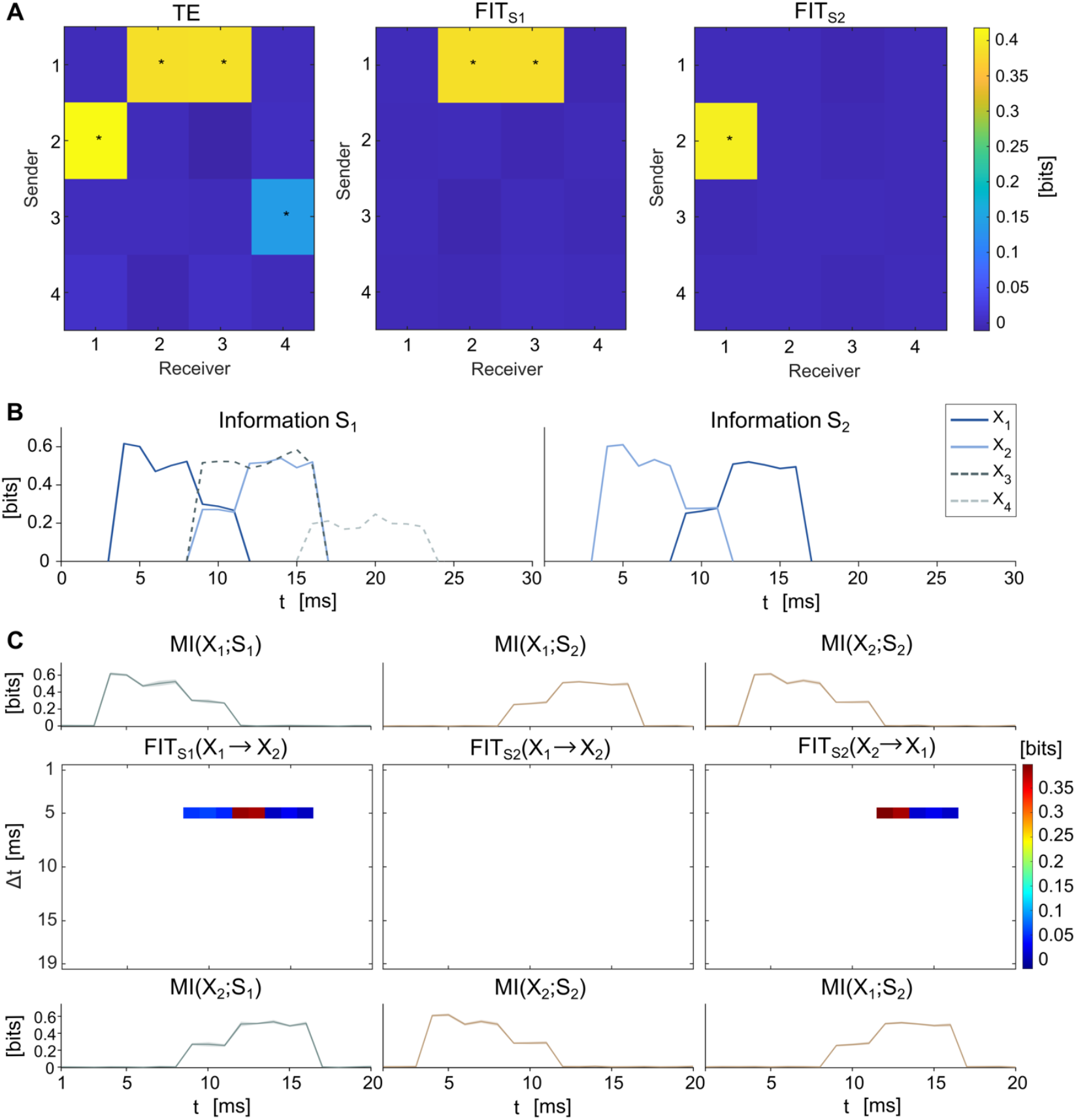
TE and FIT heatmap, temporal profiles of stimulus-feature information and time delay stimulus-feature FIT maps for X_1_ and X_2._ **A**. TE heatmap for all nodes (left), FIT about S_1_ for all nodes (middle), FIT about S_2_ for all nodes (right) for timepoint t = 12 ms with a delay of 5 ms. Significant values (p<0.01) are marked with *. **B**. Mutual Information timecourses between the nodes X and stimulus feature S_1_ and S_2_. **C**. Analysis on specific node pairs: X_1_ to X_2_ about S_1_ (left), X_1_ to X_2_ about S_2_ (middle) and X_2_ to X_1_ about S_1_ (right). **Top**. Mutual Information timecourse between the sender neural nodes and stimulus feature. **Middle**. FIT values across post-stimulus time t and delay time Δt from sender to receiver nodes about the stimulus feature. Only the time region with significant (p<0.01) stimulus information, according to a cluster permutation test with 500 null distribution samples, is plotted. **Bottom**. Mutual Information between receiver and stimulus feature. Plots show the mean across n= 10 simulations.

**Figure S6.**
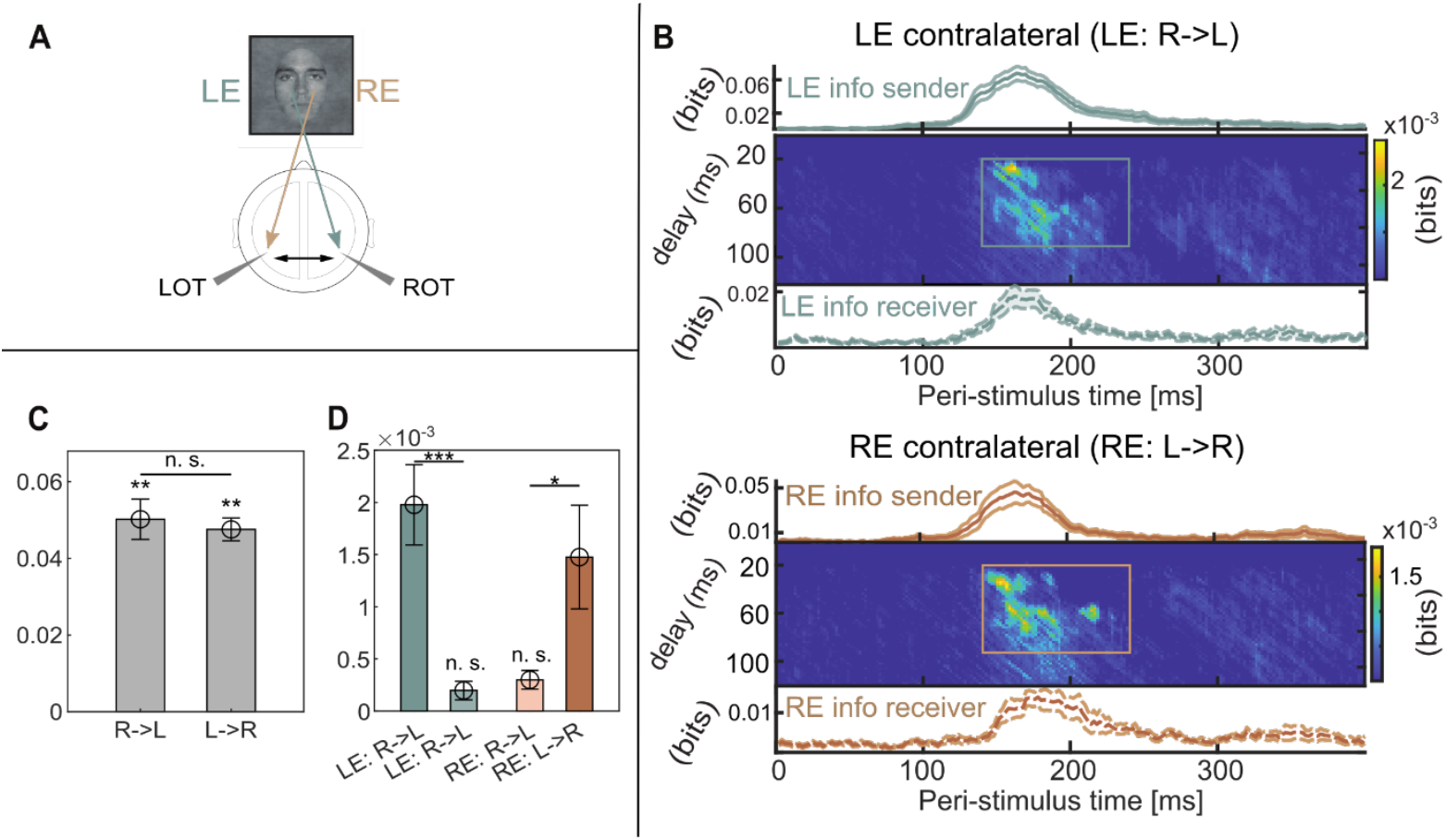
A. Schematic of the putative information flow. LOT (ROT) denote Left (Right) occipito-temporal regions. LE (RE) denotes the Left (Right) Eye visibility feature. **B**. FIT values computed across post-stimulus time *t* and delay time from sender region to receiver region about the stimulus feature. The region of the time-delay maps used to calculate the final FIT values for Fig. 4H are delimited in both heatmaps. **Top**. Mutual Information (lines) carried by the EEG in each region, and FIT (image plot) about LE contra-lateral transfer. **Bottom**. Same as Top for RE **C**. Transfer Entropy between ROT and LOT in both possible directions. **D**. FIT values computed for each stimulus (LE or RE) and direction. In this plot ROT and LOT are written as R and L, respectively. Across all panels, dots and image plots show averages. Errorbars plot SEM across participants (n=15).

